# Systematic evaluation of genome sequencing for the assessment of fetal structural anomalies

**DOI:** 10.1101/2020.08.12.248526

**Authors:** Chelsea Lowther, Elise Valkanas, Jessica L. Giordano, Harold Z. Wang, Benjamin B. Currall, Kathryn O’Keefe, Emma Pierce-Hoffman, Nehir E. Kurtas, Christopher W. Whelan, Stephanie P. Hao, Ben Weisburd, Vahid Jalili, Jack Fu, Isaac Wong, Ryan L. Collins, Xuefang Zhao, Christina A. Austin-Tse, Emily Evangelista, Gabrielle Lemire, Vimla S. Aggarwal, Diane Lucente, Laura D. Gauthier, Charlotte Tolonen, Nareh Sahakian, Christine Stevens, Joon-Yong An, Shan Dong, Mary E. Norton, Tippi MacKenzie, Bernie Devlin, Kelly Gilmore, Bradford C. Powell, Alicia Brandt, Francesco Vetrini, Michelle DiVito, Stephan J. Sanders, Daniel G. MacArthur, Jennelle C. Hodge, Anne O’Donnell-Luria, Heidi L. Rehm, Neeta L. Vora, Brynn Levy, Harrison Brand, Ronald J. Wapner, Michael E. Talkowski

**Affiliations:** Center for Genomic Medicine, Massachusetts General Hospital, Boston, MA, USA; Program in Medical and Population Genetics, The Broad Institute of M.I.T. and Harvard, Cambridge, MA, USA; Department of Neurology, Harvard Medical School, Boston, MA, USA; Program in Biological and Biomedical Sciences, Division of Medical Sciences, Harvard Medical School, Boston, MA, USA; Department of Obstetrics & Gynecology, Columbia University Medical Center, New York, NY, USA; Program in Bioinformatics and Integrative Genomics, Division of Medical Sciences, Harvard Medical School, Boston, MA, USA; Department of Pathology, Harvard Medical School, Boston, MA, USA; Department of Pathology and Cell Biology, Columbia University Medical Center, New York, NY, USA; Data Science Platform, Broad Institute of MIT and Harvard, Cambridge, MA, USA; School of Biosystem and Biomedical Science, Korea University, Seoul, South Korea; Department of Psychiatry, UCSF Weill Institute for Neuro-sciences, University of California, San Francisco, San Francisco, CA, USA; Center for Maternal-Fetal Precision Medicine, University of California, San Francisco, USA; Department of Psychiatry, University of Pittsburgh School of Medicine, Pittsburgh, PA, USA; Department of Obstetrics and Gynecology, Division of Maternal-Fetal Medicine, University of North Carolina at Chapel Hill, Chapel Hill, NC, USA; Department of Genetics, School of Medicine, University of North Carolina at Chapel Hill, Chapel Hill, NC, USA; Department of Medical and Molecular Genetics, Indiana University School of Medicine, Indianapolis, IN, USA; Division of Genetics and Genomics, Boston Children’s Hospital, Boston, MA, USA; Centre for Population Genomics, Garvan Institute of Medical Research, and University of New South Wales Sydney, Sydney, Australia; Centre for Population Genomics, Murdoch Children’s Research Institute, Melbourne, Australia

## Abstract

Current clinical guidelines recommend three genetic tests for the assessment of fetal structural anomalies: karyotype to detect microscopically-visible balanced and unbalanced chromosomal rearrangements, chromosomal microarray (CMA) to detect sub-microscopic copy number variants (CNVs), and exome sequencing (ES) to identify individual nucleotide changes in coding sequence. Advances in genome sequencing (GS) analysis suggest that it is poised to displace the sequential application of all three conventional tests to become a single diagnostic approach for the assessment of fetal structural anomalies. However, systematic benchmarking is required to assure that GS can capture the full mutational spectrum associated with fetal structural anomalies and to accurately quantify the added diagnostic yield of GS. We applied a novel GS analytic framework that included the discovery, filtration, and interpretation of nine classes of genomic variation to 7,195 individuals. We assessed the sensitivity of GS to detect diagnostic variants (pathogenic or likely pathogenic) from three standard-of-care tests using 1,612 autism spectrum disorder quartet families (ASD; n=6,448) with matched GS, ES, and CMA data, and validated these findings in 46 fetuses with a clinically reportable variant originally identified by karyotype, CMA, or ES. We then assessed the added diagnostic yield of GS in 249 trios (n=747) comprising a fetus with a structural anomaly detected by ultrasound and two unaffected parents that were pre-screened with a combination of all three standard-of-care tests. Across both cohorts, our GS analytic framework identified 98.2% of all diagnostic variants detected by standard-of-care tests, including 100% of those originally detected by CMA (n=88) and ES (n=61), as well as 78.6% (n=11/14) of the chromosomal rearrangements identified by karyotype. The diagnostic yield from GS was 7.8% across all 1,612 ASD probands, almost two-fold more than CMA (4.4%) and three-fold more than ES (3.0%). We also demonstrated that the yield of ES can approach that of GS when CNVs are captured with high sensitivity from exome data (7.4% vs. 7.8%, respectively). In 249 pre-screened fetuses with structural anomalies, GS provided an additional diagnostic yield of 0.4% beyond the combination of all three tests (karyotype, CMA, and ES). Applying our benchmarking results to existing data indicates that GS can achieve an overall diagnostic yield of 46.1% in unselected fetuses with fetal structural anomalies, providing an estimated 17.2% increase in diagnostic yield over karyotype, 14.1% over CMA, and 36.1% over ES when sequence variants are assessed, and 4.1% when CNVs are also identified from exome data. In this study we demonstrate that GS is sensitive to the detection of almost all pathogenic variation captured by karyotype, CMA, and ES, provides a superior diagnostic yield than any individual test by a wide margin, and contributes a modest increase in diagnostic yield beyond the combination of all three tests. We also outline several strategies to aid the interpretation of GS variants that are cryptic to conventional technologies, which we anticipate will be increasingly encountered as comprehensive variant identification from GS is performed. Taken together, these data suggest GS warrants consideration as a first-tier diagnostic approach for fetal structural anomalies.

## INTRODUCTION

Fetal structural anomalies are developmental defects identified by ultrasound that occur in approximately 3% of all pregnancies.^1^ The severity of structural anomalies can vary, ranging from minor defects in a single organ system to multisystem congenital anomalies associated with significant morbidity, and in some instances, mortality.^2^ Clinical genetic testing on invasively obtained DNA samples can provide critical insights into the assessment of fetal structural anomalies as the identification of a diagnostic variant in the fetus can inform medical management, prognosis, and future family planning. Current clinical guidelines recommend using gene panels as a first-tier diagnostic test when a specific genetic etiology is suspected.^3,4^ However, multiple studies have demonstrated that the phenotypic consequences for many disease-associated genes are more variable than previously recognized,^5–8^ suggesting that broad and comprehensive testing strategies are needed to maximize diagnostic sensitivity.^9^

The current standard-of-care testing for genome-wide genetic surveys involve three orthogonal and largely complementary diagnostic tests: karyotype to discover microscopically-visible balanced and unbalanced chromosomal rearrangements, chromosomal microarray (CMA) to capture sub-microscopic copy number variants (CNVs), and exome sequencing (ES) to identify single nucleotide variants (SNVs) and small insertions and deletions (indels) within the ∼2% of the genome that codes for proteins.^3,10,11^ All three tests are required to capture the full range of genetic variation currently known to contribute to fetal structural anomalies,^12–14^ which is an inefficient strategy in the prenatal setting where rapid diagnosis is critical.

Genome sequencing (GS) has the potential to identify almost all pathogenic variation captured by existing technologies in a single test.^15–17^ This technology also holds the promise of discovering novel diagnostic variants that remain cryptic to current approaches.^15,18^ Studies in rare diseases,^19,20^ and selected pediatric populations^9,21–24^ suggest that GS improves diagnostic yields over standard-of-care tests, yet systematic benchmarking of GS against individual tests performed on the same individuals from large cohorts are still lacking. These metrics are particularly important for prenatal diagnostics where a single false positive or negative result can have significant long-term consequences for the fetus and family. It is therefore imperative to assess the performance of GS against current standard-of-care approaches to determine if a shift towards recommending GS as a first-tier diagnostic test for the assessment of fetal structural anomalies is warranted.

In this study, we performed GS on 295 fetuses harboring a structural anomaly detected by ultrasound and/or a clinically relevant variant detected by standard-of-care tests. The fetuses had karyotype, CMA, and ES data available, which allowed us to identify the added diagnostic yield of GS beyond each individual test and the combination of all three tests. Comparative diagnostic yields from clinical genetic tests, such as those available for the fetuses with structural anomalies, can be biased by a variety of factors, including different test platforms, bioinformatic analyses, variant interpretation methods, assessment timepoints, sample ascertainment, and diagnostic pre-screening. Therefore, to facilitate an unbiased and robust benchmarking of our GS analytic framework on a large cohort that was not influenced by these factors, we systematically analyzed 1,612 quartet families that included a proband meeting diagnostic criteria for autism spectrum disorder (ASD), an unaffected sibling, and two unaffected parents (n=6,448 individuals) with matched GS, ES, and CMA data. Our GS analyses characterized nine different classes of genetic variation across the size and allele frequency spectrum and demonstrated high sensitivity (98.2%) against standard-of-care diagnostic tests while also identifying diagnostic variants that were missed by all three technologies. The GS analytic framework developed in this study also maintained a manageable variant review burden, which can present significant barriers to the widespread implementation of clinical GS.^25,26^ We conclude by highlighting the approaches used to interpret variants uniquely identified by GS, which will be broadly applicable to a wide variety of indications for testing.

## METHODS

### Fetal structural anomaly ascertainment and study design

We performed GS on 295 fetuses that were recruited from the Carmen and John Thain Center for Prenatal Pediatrics at Columbia University (n=193),^12,14^ the University of California San Francisco (UCSF; n=62), or the Prenatal Diagnosis Program at the University of North Carolina Chapel Hill (UNC; n=40).^27,28^ The study was approved by the Institutional Review Boards at Mass General Brigham, Columbia University, UNC, and UCSF, and written informed consent was required from both parents for study inclusion. The cohort included 249 trios identified to have a structural anomaly by ultrasound that were pre-screened with one or more standard-of-care diagnostic test (**Table S1**). The vast majority (87.6%; n=218/249) of the fetuses had no pathogenic or likely pathogenic (P/LP) variant identified by karyotype and/or CMA, and approximately one-third (35.7%; n=89/249) also had negative clinical ES (**Figure S1**). We performed the bulk of our GS benchmarking on a large cohort of 1,612 ASD quartet families (n=6,448) that had matched and unfiltered GS, CMA, and ES data available on all individuals (described below). However, we also sought to confirm these results in invasively obtained DNA samples from fetuses with structural anomalies. Therefore, we applied the GS analytic framework to 46 fetuses that were identified to carry 53 reportable variants originally detected by clinical karyotype, CMA, or ES (**Table S2** and **Figure S1**). Only the clinically relevant variants identified by each technology were available. The fetuses used for benchmarking were chosen to represent the broad types of variants known to contribute to fetal structural anomalies,^12–14^ including seven aneuploidies, eight balanced chromosomal rearrangements (BCRs), 20 CNVs, and 18 SNVs or indels (including four compound heterozygous variant pairs). Fetal DNA was obtained through an invasive diagnostic procedure and parental DNA was obtained from peripheral tissue. All DNA was extracted and sent to the Broad Institute Genomics Platform for GS, which was performed using Illumina sequencing to a mean genome coverage of 37.8 (**Tables S1** and **S2**; additional details in the **Appendix**). Sample relatedness and family structure was confirmed for all individuals using KING (**Figure S2**)^29^ and GS data was used to infer genetic sex using PLINK^30^ and depth-based chromosomal analyses (**Figure S3**; details in **Appendix**).

### Analytic framework for GS data processing and variant interpretation

We developed an analysis framework to identify P/LP variants from GS data with high sensitivity while limiting the number of variants requiring manual review per proband (**Figure 1**). The framework is organized into three components: variant discovery, variant annotation and filtration, and manual variant classification. Variant discovery identified nine different classes of genetic variation, including SNVs, indels, deletions and duplications that ranged in size from 50 base pairs to full chromosomal aneuploidies, inversions, insertions, translocations, complex rearrangements (16 different sub-classes),^31^ and short tandem repeats (STRs), using a suite of algorithms.^32–39^ All samples were jointly processed in batches following GATK Best Practices Workflows for SNV and indel discovery^38,40,41^ using Terra.^42,43^ Structural variant (SV) discovery and genotyping was performed across all samples with GATK-SV,^15,31,44^ a publicly available cloud-enabled ensemble method that leverages data from multiple SV algorithms to boost sensitivity and filters SVs to improve specificity.^15,31^ Here, we ran six SV detection algorithms^32–37,45^ on all samples and provided these data as inputs to GATK-SV, which was run in cohort mode. GATK-SV performed filtering, genotyping, breakpoint refinement, and complex variant resolution.^31^ To identify potential diagnostic STR expansion candidates, we subset the gnomAD STR catalog to 18 established disease loci that confer developmental phenotypes and ran ExpansionHunter^46^ across these loci in the fetal structural anomaly and ASD cohorts.

**Figure 1.**
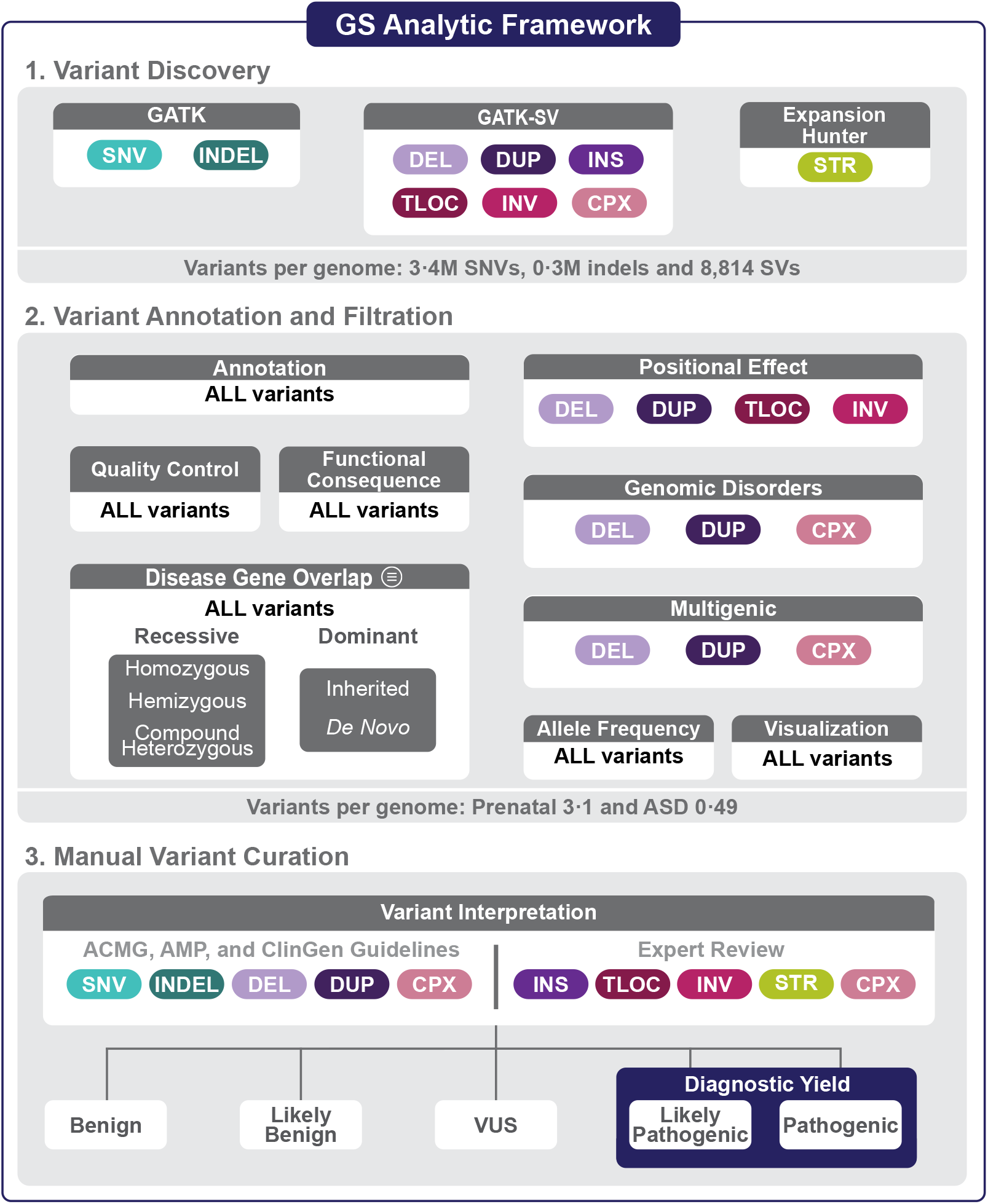
Genome sequencing analytic framework. An overview of the comprehensive framework we developed to identify diagnostic variants from GS data, which consists of three components: variant discovery, variant filtration, and manual variant curation. We considered nine different variant classes, including single nucleotide variants (SNV), small insertions and deletions (indels; below 50 base pairs), deletions (DEL) and duplications (DUP) that ranged in size from over 50 base pairs to full chromosomal aneuploidies, insertions (INS), translocations (TLOC), inversions (INV), complex rearrangements (CPX), and short tandem repeats (STRs). The filtering strategy was designed to retain P/LP variants while limiting the number of variants requiring manual review. The specific filtering criteria are described in the Appendix. All variants output by the filtering pipeline were manually curated by an expert variant review panel following existing clinical guidelines.^54–61^ All variants classified as P/LP in genes associated with the indication for testing were considered to represent the diagnostic yield of GS. VUS: variant of uncertain significance, GATK: Genome Analysis Toolkit, SV: structural variant, ClinGen: Clinical Genome Resource, ACMG: American College of Medical Genetics and Genomics, AMP: Association for Molecular Pathology.

Next, we annotated all variants for functional consequence against GENCODE v26 using ANNOVAR^47^ for SNVs and indels and svtk for SVs.^15,31^ Variants that passed our quality control filters and allele frequency thresholds were retained if they were predicted to alter a candidate disease gene or locus (see **Supplementary Methods** in the **Appendix** for details). Variants were also filtered under five genotype categories (*de novo*, rare inherited, compound heterozygous, homozygous, and hemizygous) depending on the specific mode of inheritance of the gene-disease association (dominant, recessive, or X-linked). We computationally derived our candidate disease gene list to limit the burden of up-front gene curation and to account for the phenotypic heterogeneity of the fetal structural anomaly cohort.^26,48^ We compiled a list of 2,535 genes from eight sources that are broadly associated with developmental disorders (**Table S3**; **Appendix**). SVs overlapping 64 known genomic disorder (**Table S4**) or 17 pathogenic positional effect loci (**Table S5**) were retained for further manual review.^49^ We considered STRs that exceeded a pathogenic repeat length based on literature review if they overlapped 18 STR-mediated loci that were associated with early-onset developmental disorders (**Table S6**). Finally, to remove spurious variants we visually inspected all candidate diagnostic variants output by our filtering pipeline using the Integrated Genomics Viewer for SNVs, indels, and SVs,^50^ CNView for CNVs,^51^ and REViewer for STRs.^52^**GATK**

All candidate diagnostic variants were first assessed for gene-phenotype association on a case-specific basis.^53^ If a reliable match was determined for the case in question, all variants in that gene were reviewed following the American College of Medical Genetics and Genomics (ACMG), the Association for Molecular Pathology (AMP), and the Clinical Genome (ClinGen) guidelines for SNV, indel, and CNV interpretation.^54,55^ We also incorporated recommendations for adjusting the standard clinical guidelines from the ClinGen Sequence Variant Interpretation (SVI) Working Group.^56–61^ Overall, these guidelines provide a systematic and robust method to identify variants with a 90% or greater certainty of being disease-causing.^54^ This method is reliably reproduced across laboratories^62^ and rarely results in downgrading P/LP variants over time.^63^ Candidate diagnostic variants were assessed by a variant review panel consisting of board-certified clinical geneticists, cytogeneticists, molecular geneticists, obstetricians, maternal-fetal specialists, pediatricians, genetic counselors, as well as population geneticists and bioinformaticians with expertise in SV identification and interpretation. All variants (including both variants in a compound heterozygous pair) classified as P/LP in a gene robustly associated with the case’s phenotype (*e*.*g*., the indication for testing) were considered a molecular diagnosis and were counted towards the diagnostic yield of GS.

### Systematically benchmarking GS performance

To limit the biases that can influence comparative diagnostic yields from clinical genetic tests and systematically benchmark GS against CMA and ES in a large cohort, we applied our GS analysis framework to all three technologies in 6,448 individuals across 1,612 ASD quartet families.^64^ Each quartet family comprised one proband diagnosed with ASD, one unaffected sibling, and two unaffected parents (**Table S7**). All participants or their legal guardians provided written informed consent for participation and their data were de-identified by the Simons Foundation Autism Research Initiative before sharing with qualified researchers.^64^ The quartet structure of the ASD cohort provided a unique opportunity to evaluate our GS filtering pipeline by comparing the proportion of candidate diagnostic variants identified in probands with ASD compared to their unaffected siblings. We applied the identical GS analytic framework that was implemented in the fetal anomaly cohort, except for the disease gene list used. Given that there is no universal list of clinically relevant ASD genes,^65^ we considered variants in 907 genes classified as having a ‘confirmed’ or ‘probable’ association with neurodevelopmental disorders (NDDs)^66^ and 26 genes that were enriched for rare *de novo* protein truncating variants at genome-wide thresholds in ASD (**Table S8**).^67^ We treated each ASD proband and their unaffected sibling as separate trios with both parents for filtering. To assess the potential false positive rate of the variant interpretation guidelines, we manually reviewed all variants in the ASD probands and their unaffected siblings blind to affected status (e.g., all variants were reviewed as if the child was diagnosed with ASD) and compared the fraction of P/LP variants identified between these two groups.

The ASD dataset was an ideal cohort for GS evaluation because every individual had unfiltered CMA, ES, and GS data available, which facilitated direct technology comparisons. As previously described,^45,67,68^ single nucleotide polymorphism genotyping data was generated for the entire ASD cohort using three Illumina CMA platforms and CNVs were identified from these data using PennCNV,^69^ QuantiSNPv2·3,^70^ and GNOSIS.^71^ All CNVs identified from CMA were lifted over from GRCh37/hg19 to GRCh38/hg38 for comparisons against ES and GS. The ES data were generated as part of a larger ASD sequencing initiative.^45,67^ Briefly, raw reads from all 6,448 samples were aligned to GRCh38/hg38 and SNV and indel discovery was performed using GATK v4.1.2.0.^38^ All samples were jointly genotyped following GATK Best Practices for Variant Calling.^38,40,41^ We only modified the allele balance and depth filters to accommodate the higher coverage of ES compared to GS (**Figure S4**). We also employed GATK-gCNV for exome CNV detection, a new algorithm that is specifically designed to adjust for known bias factors of exome capture and sequencing (*e*.*g*., GC content), while automatically controlling for other technical and systematic differences. The GATK-gCNV workflow is publicly available in a Terra workspace.^72^ We applied the GS analytic framework described above to the CMA and ES data and assessed the performance of GS by comparing the P/ LP variants identified by GS to those identified by CMA and ES. We also ran our GS single sample SV detection pipeline, which is described in the Appendix and is publicly available as a Terra workspace,^73^ on all ASD probands identified to have a diagnostic SV (n=77; 4.8%) to determine whether the cohort-based SV analyses provided additional value or if GS-based SV discovery can be performed on a single sample with high sensitivity and specificity against standard-of-care diagnostic tests.

## RESULTS

### Benchmarking GS in a large pediatric ASD cohort

We analyzed GS data from 1,612 ASD quartet families (n=6,448 individuals) that also had matched CMA and ES data available to directly compare the relative value of each technology (**Table S7**).^15,64,68,74^ Overall, our GS variant calling methods identified an average of 3.7M short variants (3.4M SNVs, 0.3M indels)^15^ and 8,814 SVs per genome that passed filtering criteria as well as 115,821 STR genotypes at 18 targeted disease loci across the cohort. Variant annotation and filtering reduced the number of variants requiring manual curation to an average of 0.49 variants per child (range=0-9), totaling 1,743 variants across 907 NDD-associated genes and loci in the ASD probands and unaffected siblings. We observed an enrichment of variants requiring manual review in the ASD probands compared to their unaffected siblings (0.58 mean variants per ASD proband compared to 0.39 in siblings; P=4.12×10^−14^; two-sided Wilcoxon test), suggesting that our filtering pipeline was accurately selecting for potentially pathogenic variants that should be enriched in probands. Demonstrating the power of the interpretation guidelines, this proband-sibling enrichment was further increased following manual variant curation, which identified 128 P/LP variants in 126 ASD probands (7.8%; 95% CI 6.5-9.1) compared to 17 unaffected siblings (1.1%; 95% CI 0.6-1.6; odds ratio [OR]=7.9; 95% CI=4.7-14.1; P=2.2×10^−16^; Fisher’s exact test) (**Figure 2** and **Table S9**) and differed significantly across IQ subgroups in ASD probands (**Figure S5**).

**Figure 2.**
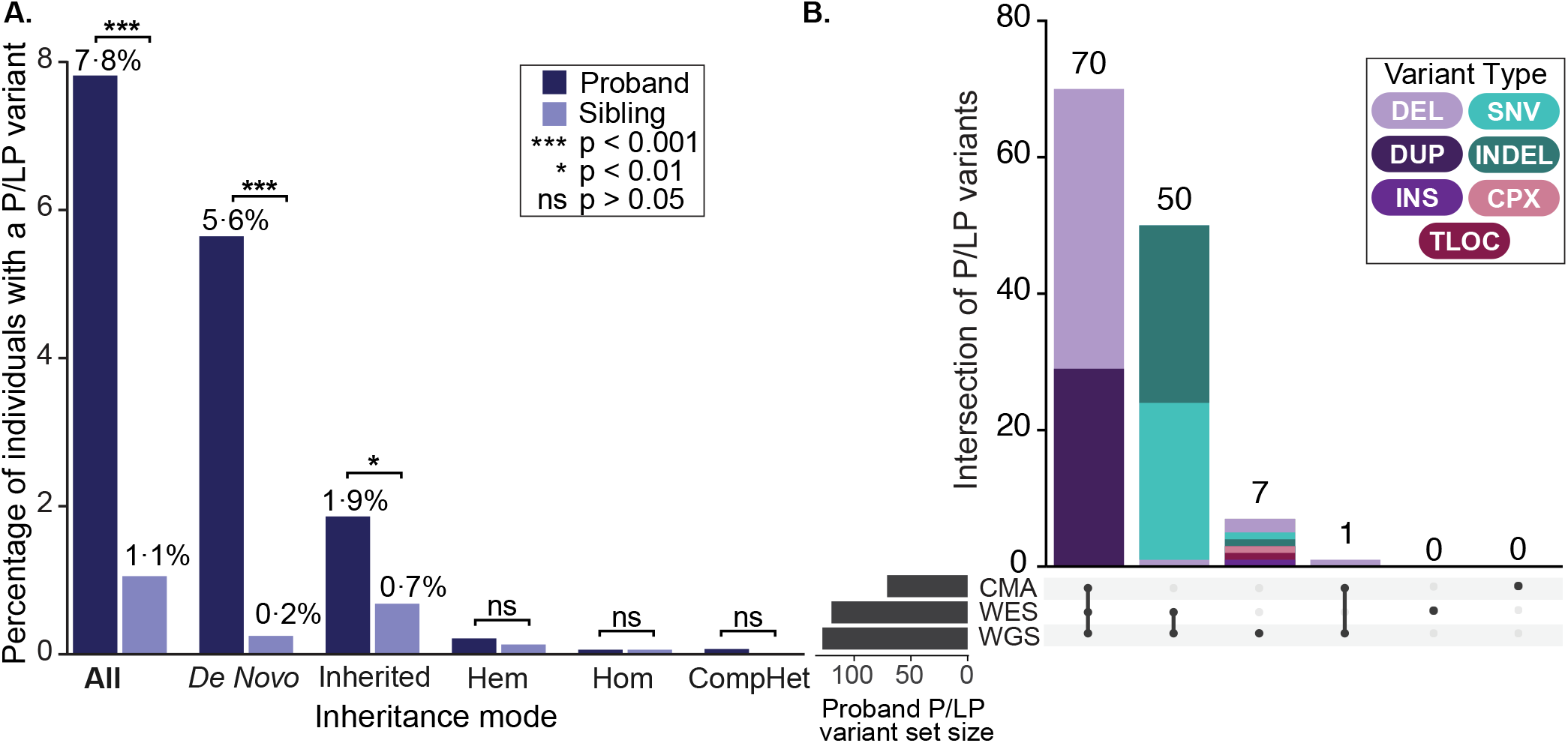
Benchmarking the performance of GS in ASD probands and unaffected siblings. **(A)** The fraction of ASD probands and unaffected siblings identified to carry a P/LP variant by GS subset by inheritance category. The denominator used for all categories was 1,612 except for hemizygous variants where only males were considered (n=1,440 male probands and 755 male siblings). **(B)** The total number of P/LP variants (n=128) detected by each technology (GS, CMA, and ES) in ASD probands.

We benchmarked the diagnostic performance of GS against standard-of-care tests by applying the equivalent GS framework to the CMA and ES data, with minor modifications to accommodate each data type (**Figure S4**; **Appendix**). Overall, GS identified a diagnostic variant in almost two-fold more probands than CMA (n=126 vs. n=71; 4.4%; OR=1.8; 95% CI 1.3-2.5; P=6.5×10^−5^) and almost three-fold more than ES (n=126 vs. n=49; 3.0%; OR=2.7; 95% CI 1.9-3.9; P=1.98×10^−10^) (**Figure 2**). However, when we used a new method to capture CNVs from ES data (GATK-gCNV),^72^ the overall diagnostic yield of ES approached that of GS (7.4% vs. 7.8%, respectively), but still did not capture all known P/LP CNVs. For example, a 504kb deletion overlapping the first exon of *NRXN1* was present in the raw ES CNV calls but did not pass our ES filtering, which required >3 exons to be overlapped. This suggests strategies for clinical exome CNV calling should consider relaxing filtering for well-known disease genes, particularly for genes where CNVs are a known mechanism of disease.^75^ Conversely, GS captured 100% of the P/LP variants identified by CMA (n=71) and ES (n=118; n=70 variants were identified by both CMA and ES). Similarly, the single sample GATK-SV pipeline detected all 77 diagnostic SVs present in ASD probands (**Table S9**), indicating that diagnostic laboratories can use the normalization procedures provided in the single-sample GATK-SV algorithm and do not need to accumulate large cohorts of GS data to perform comprehensive SV discovery on individual samples.

GS identified an additional diagnostic variant in seven ASD probands (**Figure 2**), representing a 0.4% increase in diagnostic yield beyond the combination of both CMA and ES. The GS-unique variants included one SNV and one indel, a *de novo* stopgain in *ANKRD11* and a 44bp *de novo* frameshift insertion in *SMARCA4*, and five SVs, two in-frame single exon deletions in *RERE* and *RORA*,^76,77^ a reciprocal translocation disrupting *GRIN2B*, a SVA retrotransposon insertion in *DMD*, and a 47.2Mb complex SV involving chromosome 1 that comprised four deletions, an inversion, and an inverted insertional translocation. The *ANKRD11* stopgain was in an exon with no ES coverage and the *SMARCA4* insertion was within 30bp of an intron-exon boundary and was not detected in the ES read evidence (**Figure S6**). In contrast to the single exon *NRXN1* deletion described above, the smaller *RERE* (5.6kb) and *RORA* (0.5kb) deletions identified by GS were not detectable in the raw ES data, suggesting that gene-specific CNV filtering will not be able to re-capture all single-exon deletions of clinical relevance. As expected, CMA and WES were unable to detect the diagnostic balanced translocation. Similarly, while CMA and ES detected the four *de novo* deletions involved in the complex SV, they were unable to identify the inversions that would have linked the deletions into a single event. Given the number of rearranged segments and their relative size, cytogenetics methods alone would have also been unable to fully resolve the complex event. Indeed, GS identifies complexity that is cryptic to karyotype in approximately one out of every four cases harboring an apparently BCR.^78^ Taken together, these data demonstrate that GS outperforms both CMA and ES, capturing all P/LP variants from these two technologies, while providing a modest increase in diagnostic yield beyond the combination of both diagnostic tests.

### Application of GS for the assessment of fetal structural anomalies

After systematically benchmarking the performance of our GS analytic framework, we applied it to 249 fetus-parent trios that were pre-screened with karyotype, CMA, and/or ES (**Figure S1**). The structural anomalies impacted a wide range of organ systems, with 36.1% (n=90/249) of the cohort having multisystem involvement (**Figure 3** and **Table S1**). GS identified 816 candidate variants requiring manual review, resulting in an average of 3.1 variants per fetus (median=3.0, range=0-21). The increased number of variants output by our GS filtering in the fetuses compared to ASD probands reflects the greater number of genes considered: n=2,535 genes for the fetal cohort versus n=907 genes for the ASD cohort. Manual variant curation identified twenty-one P/LP variants in nineteen fetuses with a structural anomaly, suggesting that the added diagnostic yield of GS beyond first-tier diagnostic tests (CMA and karyotype) is 7.6% (**Table S10**). Interestingly, the GS yield is relatively similar to previous ES studies of fetal structural anomalies (**Figure S7**)^13,14^ despite the fact that over one-third of our cohort was depleted for diagnostic variants that are identifiable by ES.

**Figure 3.**
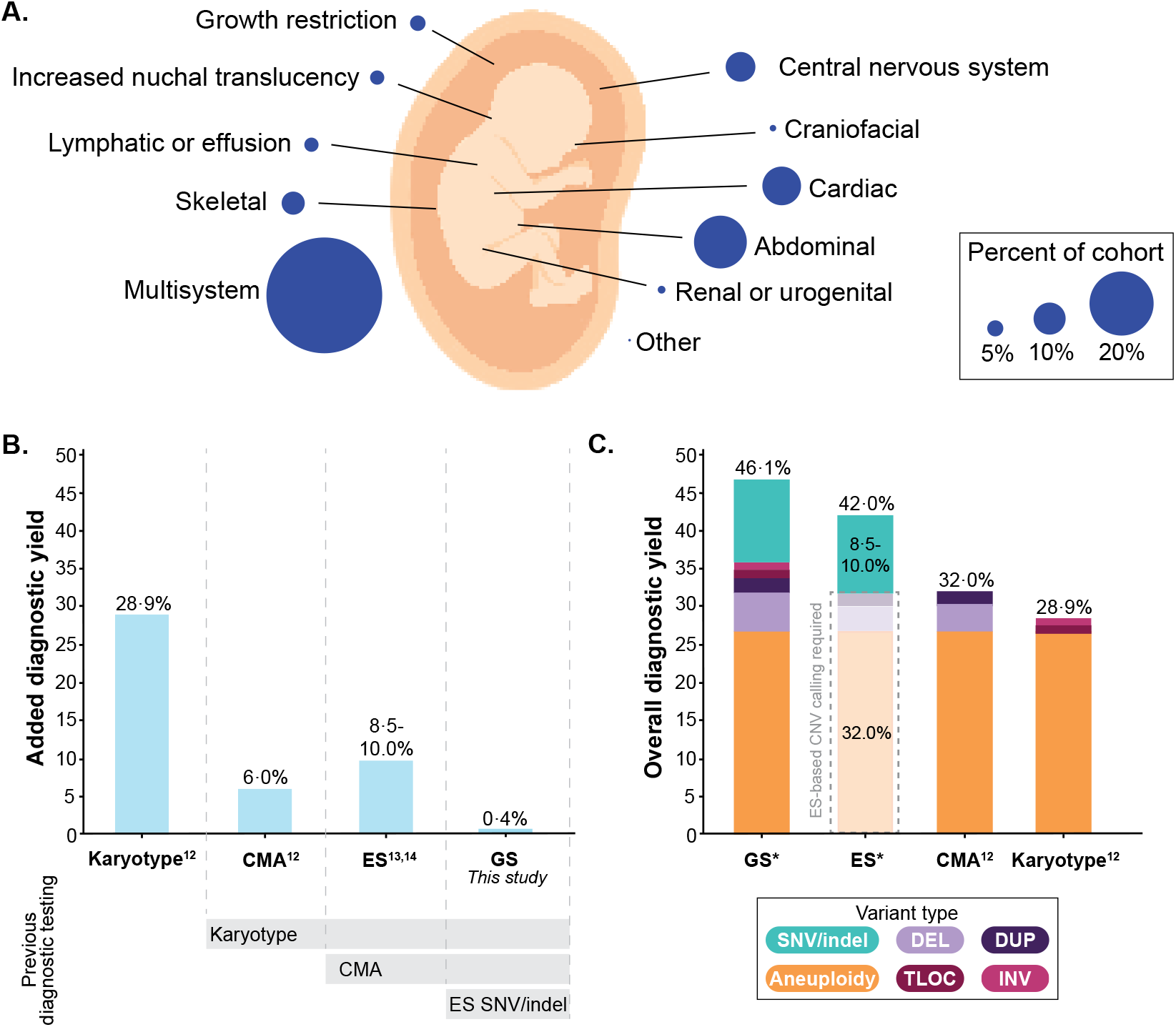
Overview of fetuses with structural anomalies and diagnostic yields across technologies. **(A)** The phenotypic breakdown of 249 trio fetuses identified to have a structural anomaly detected by ultrasound included in this study. The fetuses were pre-screened with a combination of standard-of-care diagnostic tests (see Methods for details). Fetuses with anomalies impacting more than one body system were counted as having multisystem abnormalities. The remaining categories represent fetuses with isolated structural anomalies. **(B)** The added diagnostic yield for each sequencing technology when applied serially to pre-screened cases. Each technology is assessed in a cohort that was depleted for diagnostic variants detected by the preceding technology. Yields for karyotype and CMA were taken from Wapner et al.^12^ and the diagnostic range for ES from Petrovski et al.^14^ and Lord et al.^13^ **(C)** The anticipated overall diagnostic yield provided by each diagnostic test if they were applied to a cohort of unselected fetuses with structural anomalies. *The yields for ES and GS were predicted based on data from this study as well as previously published work. The dashed grey box surrounding the ES bar indicates the diagnostic yield that could be captured if ES-based CNV methods are applied.^12,14^ Each bar is colored based on the fraction of diagnoses provided by each variant class. CMA: chromosomal microarray, ES: exome sequencing, GS: genome sequencing, SNV: single nucleotide variant, indel: small insertion and deletion, CNV: copy number variant, DEL: deletion, DUP: duplication, TLOC: translocation, INV: inversion.

The P/LP variants identified by GS included 15 SNVs, two indels, two independent *de novo* SVs in *MED13L* in unrelated fetuses, a compound heterozygous variant pair comprising a missense variant in *trans* with a 143 kb intragenic exonic duplication in *DYNC2H1*, and a maternal uniparental disomy (UPD) event involving chromosome 20 (**Table S10**). No P/LP STR expansions across the 18 disease-associated loci were identified. The benchmarking results from our ASD cohort predict that the majority (n=20; 95.2%) of these P/LP variants could be identified by CMA or ES had it been performed using contemporary platforms, variant calling algorithms, and analysis pipelines. Interestingly, due to the specificity of the gene-phenotype association for the missense variant in *DYNC2H1*,^79^ the diagnostic laboratory manually reviewed the ES read depth profile across this gene, identified the duplication, and confirmed the event with fluorescence *in situ* hybridization.^27^ The most conservative estimate therefore suggests that GS uniquely identified one diagnostic variant, a single exon deletion in *MED13L* (1.3kb in size) resulting in at least a 0.4% increase in diagnostic yield beyond the combination of three independent tests (karyotype, CMA, and ES) in fetal structural anomalies if state-of-the-art methods are universally applied (**Figure 3**).

We next wanted to confirm the GS benchmarking results from the ASD quartet families using DNA from invasively obtained prenatal samples as well as assess the performance of GS to detect chromosomal rearrangements routinely identified by karyotype, a standard-of-care diagnostic test for fetal structural anomalies that was not widely available for the ASD cohort. We chose 46 fetuses that carried 53 reportable variants that broadly represented the types of genomic variation known to contribute to fetal structural anomalies (**Figure S1** and **Table S2**).^12–14^ Overall, GS captured 94.3% (n=50/53) of the clinically reportable variants, including 100% of aneuploidies (n=7), CNVs (n=20), and SNVs/indels (n=12), as well as five of the eight balanced chromosomal rearrangements (BCRs; **Figure S1**). The three variants not captured were BCRs that involved telomeric or centromeric regions based on the reported karyotype. These data were consistent with our prior studies from long-insert GS,^49,78,80,81^ where GS discovered all BCR breakpoints located in unique sequence, but was unable to capture breakpoints in highly-repetitive regions known to be inaccessible to short-read sequencing.^82^ Given the high sensitivity of GS to detect all clinically relevant variants from standard-of-care CMA and ES diagnostic tests, and the majority of BCRs, we sought to estimate the potential diagnostic yield of GS in a cohort of unselected fetal structural anomalies. To accomplish this, we extrapolated our benchmarking data to two previously published studies describing the yield of karyotype, CMA, and ES in unselected fetal structural anomalies^12,14^ and estimate that GS can provide an overall diagnostic yield of approximately 46.1% (**Figure 3**). This suggests that GS significantly outperforms each individual standard-of-care test by a wide margin, providing an estimated 17.2% increase in diagnostic yield over karyotype, 14.1% over CMA, and 38.3% over ES when only SNVs and indels are considered. We estimate that the application of a CNV calling algorithm to the ES data would result in GS providing a 4.1% increase in diagnostic yield over ES when performed on unselected fetuses with structural anomalies (**Figure 3**).

### Classification of SVs unique to GS

The eight diagnostic variants uniquely identified by GS in the ASD and fetal structural anomaly cohorts included SNVs and indels in exons not captured by ES (n=2), SVs below the resolution and/or inaccessible to karyotype, CMA, or ES (n=4), and SVs for which the resolution of GS resulted in a medically relevant change in classification from VUS to LP/P (**Figure 4**).^63^ Given the relatively limited number of cases for which GS diagnostics has been performed to date, and the myriad functional consequences that can result from CNVs and other classes of SVs that should now be routinely discovered in each human genome, SVs continue to present a considerable interpretative challenge. To highlight this complexity and support reproducibility for SV classification, we outline how the existing clinical interpretation guidelines^55,58^ were used to address this diverse class of variation that collectively accounts for >25% of all predicted loss-of-function (LoF) variants in each human genome.^31^

**Figure 4.**
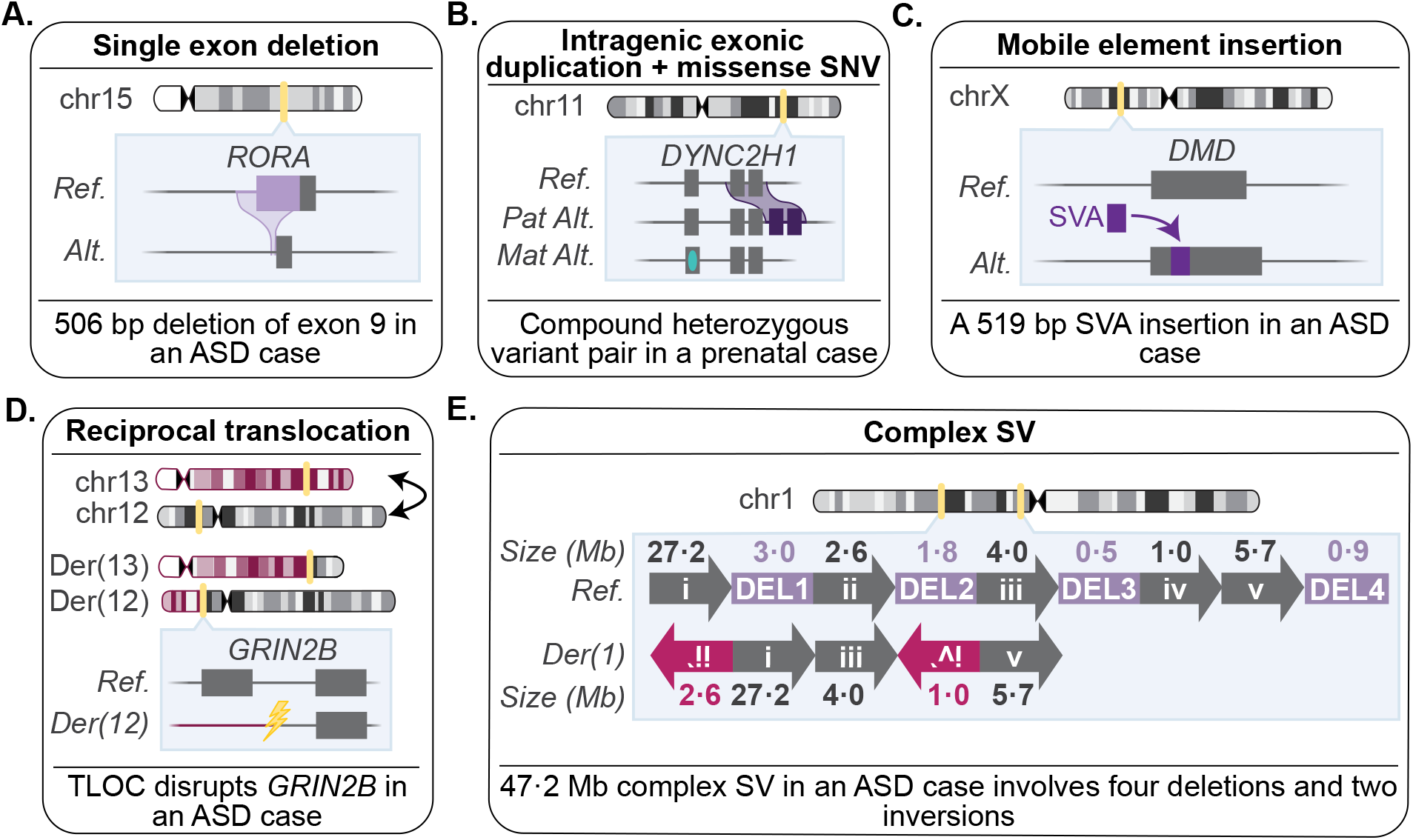
Diagnostic structural variants unique to genome sequencing. **(A)** A 506bp single exon in-frame deletion in *RORA* in an ASD proband. **(B)** A compound heterozygous missense variant in *trans* with an intragenic exonic duplication in *DYNC2H1* in a fetus with short-rib thoracic dysplasia. **(C)** A SVA retrotransposon insertion disrupting *DMD* in an ASD proband. **(D)** A balanced reciprocal translocation between chromosomes 12 and 13 in an ASD proband that directly disrupts *GRIN2B*. **(E)** Linear representation of a *de novo* complex SV impacting chromosome 1 in an ASD proband. Each rearranged segment of DNA in the derivative chromosome is depicted by a unique roman numeral (i-v), while the four deleted segments of DNA are outlined in purple and sequentially numbered DEL 1-4 (6.3 Mb total deleted). Arrows are not drawn to scale, and inverted segments are denoted by a reverse orientation of arrows. Genomic coordinates are provided in Table S11.

Three of the four GS-unique diagnostic SVs included small *de novo* CNVs below the resolution of CMA and ES. In some cases, predicting which of these variants were LoF was clear, as in a 1,369 bp *de novo* deletion in exon 14 of *MED13L* that caused a frameshift in a fetus with a lymphatic malformation and pleural effusion. In contrast, intragenic deletions like the two in-frame single exon *de novo* deletions in *RERE* and *RORA* in unrelated ASD probands were more challenging to interpret.

We used the ClinGen SVI Working Group recommendations^58^ for appropriately weighting variants previously assumed to truncate or eliminate gene products that do not ultimately result in nonsense mediated decay. Both in-frame *de novo* deletions disrupted less than 10% of their respective gene’s protein-coding sequence, did not disrupt any known functional domains in either protein, and no additional cases with deletions small enough to associate these particular exons with disease were found in the literature or open-access variant databases. However, there was a complete absence of protein-coding deletions in *RERE* and *RORA* among 14,891 population controls from the genome aggregation database (gnomAD),^31^ which applied similar SV detection methods as our study, which we considered sufficient evidence to classify these two deletions as LP. To improve the quantitative approach to classifying SVs only accessible to GS, additional studies will be required to fully understand the factors that influence the likelihood that small in-frame CNVs are disease-causing, particularly for genes where exon-resolution functional characterization has yet to be performed and cannot be used to inform variant interpretation.

In two ASD cases, GS identified a balanced translocation and a complex SV that may have been potentially identifiable by karyotype but could only be classified as diagnostic using the additional information uniquely provided by GS. For example, while reciprocal translocations identified by karyotype are routinely reported back to families, very little can be said about their contribution to the indication for testing because the precise location of their breakpoints remains unknown.^49,78,83^ Indeed, our previous work has demonstrated that GS revises the location of cytogenetically visible BCRs by one or more cytogenetic band in over 93% of cases.^78^ This suggests GS provides BCR breakpoint resolution that negates the need to generate unique primers each time a novel BCR is identified, which is the process many diagnostic laboratories offering karyotype follow. The translocation breakpoint identified in an ASD proband is in intron two of *GRIN2B*. In contrast to other null variants,^54,58^ BCR breakpoints located in introns as well as exons can result in LoF, as long as alternative splicing, transcript diversity, and proximity to 3’ exons are taken into consideration.^54,58^

The GS data also discovered a *de novo* 47.2 Mb complex SV in an ASD proband that involved six breakpoints: two inversions and four deletions that disrupted 42 protein-coding genes in total, none of which have been previously associated with ASD or are supported by case evidence in the literature (**Figure 4** and **Table S11**). Current guidelines recommend the assessment of individual CNVs involved in a complex SV rather than evaluating the overall rearrangement to arrive at a final diagnostic classification.^55^ Strictly following the guidelines would result in a VUS classification because the largest deletion only overlapped 18 genes, which did not meet the minimum threshold of 35 genes to be classified as LP.^55W^ However, GS data enabled the discovery that all four deletions were part of the same rearrangement, which provided strong rationale to apply the gene count threshold to the entire complex SV, resulting in a classification of LP. We note that complex SVs identified by GS were not included in the original analyses that defined the gene-count thresholds,^55^ suggesting that future studies could investigate whether complex SVs follow similar gene disruption patterns as canonical deletions and duplications. In summary, these data provide specific examples of the challenges and solutions for interpreting GS unique variants and highlight the type of variants that will be increasingly encountered as comprehensive variant identification from clinical GS becomes more widespread across a wide variety of phenotypes.

## DISCUSSION

Since the advent of massively parallel sequencing technologies, the application of clinical GS has represented an enticing approach to ascertain almost all pathogenic variation in a single diagnostic test. With the exception of a few small prenatal studies of fetal structural anomalies,^84–87^ the overwhelming majority of research examining the clinical utility of GS has been performed in the pediatric setting. For example, multiple studies have demonstrated that GS provides a higher diagnostic yield than standard-of-care tests across a wide range of pediatric phenotypes,^9,21–23,88,89^ and recent guidelines now recommend GS as a first-tier diagnostic test for the assessment of intellectual disability and/or congenital anomalies.^90^ However, these pediatric cohorts frequently have disparate standard-of-care tests available,^9,21,88,91^ precluding systematic benchmarking of GS against any individual test. Further, existing clinical GS studies have either not considered SVs^88,89,92^ or relied on a small number of algorithms for their detection.^9,22,23,91^ This places an unnecessary technical constraint on the diagnostic value of GS and represents a critical limitation for prenatal diagnostic testing where the contribution of SVs to phenotypes such as fetal structural anomalies is significant.^12^ We demonstrate here that these limitations can be circumvented using a comprehensive GS framework to capture, filter, and interpret a broad spectrum of variant classes without significantly increasing the burden of manual variant curation.^26^ We accomplished this by systematically benchmarking the performance of our GS analysis framework against CMA and ES in a large pediatric cohort and then applying that framework to assess the diagnostic utility of GS as a first-tier diagnostic test for fetuses with structural anomalies.

Given the critical need to balance sensitivity and specificity for genetic testing in the prenatal setting, we first sought to conduct an extensive benchmarking study that included the simultaneous analysis of three technologies (GS, ES, and CMA) in a large cohort of 1,612 ASD quartet families. This allowed us to perform direct technology comparisons on the same individuals that were not influenced by different testing platforms, assessment timepoints,^93^ bioinformatic analyses, or inconsistent application of interpretation guidelines.^62^ We demonstrated that GS captures all diagnostic variants identified by CMA and ES, and provides a molecular diagnosis for almost two-fold more ASD probands than either technology alone. We also illustrate that the diagnostic yield of ES can approach that of GS if CNV discovery is performed on the exome data with a sufficiently sensitive tool.^94–96^ To confirm these results in invasively obtained DNA samples from fetuses, we applied our GS analysis framework to 46 fetuses that had a reportable variant identified by clinical karyotype, CMA, or ES. Based on these and existing data,^12,14^ we estimate that the potential diagnostic yield of GS in unscreened fetuses with structural anomalies to be 46.1%, suggesting that GS significantly outperforms each conventional test, proving an estimated 17.2% increase in diagnostic yield over karyotype, 14.1% over CMA, and 36.1% over ES when only SNVs and indels are considered. We note, however, that the exact yield of GS will be highly dependent on the phenotype distribution of the cohort being studied.^13,14,97^ Together, these data strongly argue for GS to displace the serial application of karyotype, CMA, and ES for the assessment of fetal structural anomalies, provided analysis and interpretation are sufficiently optimized to identify and interpret all classes of variation.

Overall, GS uniquely identified eight P/LP variants across ASD probands and fetuses with structural anomalies, representing an added diagnostic yield of 0.4% in both cohorts. This estimate is significantly lower than existing pediatric studies that found GS could provide a 9-28% increase in diagnostic yield when compared to a wide variety of standard-of-care diagnostic tests.^9,22,90,98^ While phenotype, ascertainment, and clinical context are expected to impact GS yields, our study demonstrates the importance of performing systematic benchmarking of the same tests on multiple individuals combined with comprehensive variant discovery from all technologies to avoid inflating the added diagnostic yield that GS currently provides. Our study revealed that most diagnostic GS-unique variants included SVs that were inaccessible to existing standard-of-care diagnostic tests or were only determined to be diagnostic based on information that was uniquely provided by GS. These included BCRs, complex SVs, single exon in-frame deletions, and mobile element insertions. We highlight several ways to use the existing clinical guidelines^55,58^ to accommodate the interpretation of these variants, which we expect to be increasingly encountered as clinical GS becomes more widespread. While the added diagnostic yield of GS is expected to differ across phenotypes, the unbiased assessment performed herein using large samples, state-of-the-art molecular and computational methods, and updated interpretation guidelines should temper enthusiasm regarding immediate significant increases in interpretable pathogenic variation from GS. Absent improvements in variant annotation, especially for noncoding variation, advances in genomics technologies will continue to only provide incremental increases in diagnostic yield.

In addition to the performance and diagnostic yield of GS, there are technical, logistical, and economic considerations for laboratories and clinicians when deciding to implement a new diagnostic test such as GS. Among these, technical capacity and timely return-of-results is paramount in the prenatal setting. While assessing turn-around-time was beyond the scope of this study, rapid GS (ranging from 26 hours to 3.2 days for analysis completion)^88,99^ has been demonstrated in the pediatric setting for the assessment of critically ill infants, where, similar to the prenatal diagnostics, time to diagnosis can have a significant impact on medical management and clinical outcomes. Further, the cost of clinical GS has been shown to be lower than existing standard-of-care diagnostic tests for individuals with a developmental disorder and/or congenital anomaly^100^ and rapid GS has been shown to reduce the cost of hospitalization for children admitted to neonatal or pediatric intensive care units.^101^ Taken together, these data suggest that the benefits of GS are likely to extend beyond rapid return-of-results and improved diagnostic yield. A single comprehensive test like GS will also preserve precious fetal DNA. Perhaps the most important factor when considering implementing GS for prenatal genetic diagnosis is the accuracy of variant interpretation. The return of variants that are unrelated to the indication for testing or associated with benign outcomes can have significant negative consequences for families. It is therefore notable that our phenotype-blinded variant interpretation resulted in just 17 variants meeting criteria for a P/LP classification in 1,612 unaffected ASD siblings. Importantly, 71% of these P/ LP variants in siblings included CNVs associated with reduced penetrance (Table S9), which are known to be a challenge for genetic counseling and are already encountered by clinicians during routine prenatal CMA testing. These data indicate that the application of rigorous variant interpretation guidelines should result in few fetuses being misdiagnosed.

In conclusion, these studies demonstrate the potential for GS to displace a series of standard-of-care diagnostic tests that individually identify only a small portion of the genomic variant spectrum associated with fetal structural anomalies. These analyses represent the largest effort to benchmark the performance of GS to date. The scale of such benchmarking is critical, as these analyses focus on rare variation that span an array of mutational mechanisms but are not frequently observed in the general population or in small cohorts of fetuses with structural anomalies.^84–87^ We developed a framework to systematically detect pathogenic SNVs, indels, CNVs, balanced SVs, complex SVs, and STRs at base-pair resolution and applied the most up-to-date interpretation guidelines from ACMG/AMP and ClinGen.^54–61^ We demonstrate that GS is unlikely to significantly increase the diagnostic yield in fetal structural anomalies without improvements in variant annotation and interpretation, particularly for noncoding variation as we were only able to consider a small number of noncoding disease-associated loci. Further, some discrete phenotypes will continue to require specialized assays (*e*.*g*., methylation tests, microsatellite analysis) for variants not accessible to any NGS technology. We expect that, similar to existing variant classes, GS-unique variants will also benefit from the accumulation of classified variants in open-access databases such as ClinVar.^103^ Overall, these data suggest that GS can effectively displace karyotype, CMA, and ES as a single diagnostic test for the assessment of fetal structural anomalies and will provide a marginal, but important, increase in diagnostic yield beyond the combination of all three standard-of-care diagnostic tests.

## ACKNOWLEDGEMENTS

We thank the families and clinicians from the Columbia University Carmen and John Thain Center for Prenatal Pediatrics, the University of North Carolina (UNC) Chapel Hill Prenatal Diagnosis Program, the University of California San Francisco Prenatal Diagnostic Center, and the Simons Simplex Collection for their participation. This study was supported by resources from the National Institutes of Health (NIH): HD081256, HD099547, and MH115957 (awarded to M.E.T.), HD088742 (awarded to N.V.), UM1HG008900 (awarded to A.O.L., H.R., and M.E.T), and the Simons Foundation for Autism Research Initiative (SFARI #573206 awarded to M.E.T.). C.L. was supported by a postdoctoral fellowship from the Canadian Institutes of Health Research. E.V. was supported by NINDS F31NS113414. R.L.C. was supported by NHGRI T32HG002295 and NSF GRFP #2017240332. H.B. was supported by NIDCR K99DE026824. J-Y.A. was supported by NRF-2020R1C1C1003426 and NRF-2017M3C7A1026959.

## AUTHOR CONTRIBUTIONS

Study design: B.L., H.B., D.G.M., R.W., and M.E.T. Family recruitment and sample collection: J.L.G., V.S.A., D.L., M.E.N., T.M., K.G., B.P., A.B., M.D., N.L.V., B.L., and R.W. Sample library preparation: B.B.C. and K.O’K. Computational analysis: C.L., E.V., H.Z.W., E.P.H., N.K., C.W.W., S.P.H., B.W., V.J., J.F., R.L.C., X.Z., L.D.G., C.T., N.S., J-Y.A., S.D., B.D., D.B.G., S.J.S., D.G.M., and H.B. Manual variant curation: C.L., E.V., J.L.G., C.A.A-T., E.E., G.L., K.G., F.V., J.C.H., A.H.O’D-L., H.L.R., N.L.V., B.L., and R.W. Verified the underlying data for these analyses: C.L., E.V., H.B., and M.E.T. Wrote the manuscript and generated the figures: C.L., E.V., H.B., and M.E.T. All authors reviewed the manuscript. C.L. and E.V. contributed equally to this study.

## DATA SHARING

The genomic and phenotype data for the ASD families can be accessed through SFARIbase with permission from the Simons Foundation Autism Research Initiative. The CRAM files from the trios with fetal structural anomalies will be made available through AnVIL at publication.

