## Supplementary Appendix for "Systematic evaluation of genome sequencing for the assessment of fetal structural anomalies"

Lowther C*, Valkanas E* *et al.*
*Full list of authors is available in the main text*

#### Table of Contents

**SUPPLEMENTARY METHODS** 1
- Participant ascertainment 1
- Genome sequencing and sample-level QC 1
- Genome sequencing analysis framework 1
  - 1.0. Variant discovery 2
    - 1.1. Sequence variants (GATK) 2
    - 1.2. Structural Variants (GATK-SV) 2
    - 1.3. Short tandem repeats (Expansion Hunter) 3
  - 2.0. Variant filtering 3
    - 2.1. Variant QC 3
    - 2.2. Variant functional consequence 3
    - 2.3. Disease genes and genomic regions 4
    - 2.4. Inheritance 6
    - 2.5. Allele frequency 7
  - 3.0. Variant interpretation 7
- Single sample SV pipeline 7
- Benchmarking the performance of GS against conventional tests 7
- Filtering CMA data 7
- Filtering exome sequencing data 9

**SUPPLEMENTARY FIGURES** 9
- Figure S1. Prenatal study design 9
- Figure S2. Confirmation of sample relatedness using kinship values 10
- Figure S3. Sample sex QC 11
- Figure S4. Modified exome sequencing depth and allele balance thresholds 12
- Figure S5. Fraction of P/LP variants across IQ subgroups in ASD probands 13
- Figure S6. Two pathogenic sequence variants unique to GS in ASD probands 14
- Figure S7. Diagnostic yield of ES and GS across organ systems impacted by fetal structural anomalies 15

**REFERENCES** 16

### SUPPLEMENTARY METHODS

#### Participant ascertainment

We ascertained 295 fetuses with a structural anomaly detected by ultrasound for inclusion in this study. Recruitment and phenotyping protocols for the fetuses have been previously published.[1–3] We also included 1,612 deeply phenotyped quartet families ascertained from the Simons Simplex Collection in this study.[4–6] As previously described,[7] each family included two unaffected parents, one unaffected sibling, and an affected proband with autism spectrum disorder (ASD). All affected probands underwent a battery of diagnostic tests, including the Autism Diagnostic Observation Schedule (ADOS) and the Autism Diagnostic Interview-Revised (ADI-R) to confirm the ASD diagnosis, as well as detailed evaluations of intellectual/cognitive functioning, adaptive behavior, physical/dysmorphic features, developmental milestones, medical comorbidities, and family history. Multiple assessment measures were performed in the parents and siblings to exclude the presence of ASD symptoms (https://www.sfari.org/resources/ssc-instruments/).[4] All 7,195 individuals underwent paired-end genome sequencing (GS) to a mean target coverage of 30X (see Table S1, S2, and S7 for specific sequencing metrics).

#### Genome sequencing and sample-level QC

To confirm sample relatedness we performed a kinship inference analysis with KING[8] (http://people.virginia.edu/~wc9c/KING) using the GS data after restricting to single nucleotide polymorphisms (SNPs) with an alternate allele frequency (AF) >5% in gnomAD (Figure S2).[9] In parallel, we also predicted genetic sex using two independent approaches: first, we used PLINK to infer sex based on sex chromosome genotypes.[10] Second, we used GATK-SV to calculate copy number estimates for each chromosome per sample, which permitted inference of genetic sex as well as the identification of chromosomal aneuploidies. We compared the predicted sex for each individual between both methods and observed high concordance (Figure S3). Using the relatedness and sex results we resolved discrepancies in family structures deviating from the expected relatedness metrics for parent-child (IBS0 ≤ 0.005 and kinship coefficient > 0·2) and sibling relationships (IBS0 > 0·005 and kinship coefficient > 0·2; Figure S2).

We also confirmed cross-technology sample relatedness to assure comparisons were performed on the same 6,448 individuals from the 1,612 ASD quartet families. This was accomplished by restricting the exome sequencing (ES) and GS VCFs to high-quality common SNPs from Purcell et al. 2014[11] that were lifted over to GRCh38/hg38 and limited to 5,862 SNPs common to both ES and GS. Samples were renamed based on their technology of origin and were merged into a single cross-technology ASD VCF for relatedness analysis with KING.[8] ES and GS samples with a kinship coefficient > 0·45 were considered to be identical samples. Finally, we used the confirmed sample metadata from the GS and ES comparisons to identify matching CMA data.[12]

#### Genome sequencing analysis framework

We developed a GS analytic framework to discover, filter, and interpret nine different classes of variation that are described in detail below and also summarized in Figure 1. The aim of this pipeline was to retain as many pathogenic or likely pathogenic (P/LP) variants as possible while reducing the total number of variants requiring manual review. The same GS analysis pipeline was applied to both the ASD and fetal anomaly cohorts with only modifications to the phenotype-specific gene list used.

*1.0. Variant discovery*

1.1. Sequence variants (GATK)

As previously described,[5,6] the ASD GS data was generated from PCR-free libraries and processed using the Center for Common Disease Genomics functional equivalence pipelines (https://github.com/CCDG/Pipeline-Standardization) and following the Genome Analysis Toolkit (GATK) Best Practices Workflows for sequence variant, single nucleotide variant (SNV) and small insertion/deletion (indel), discovery.[13–15] Briefly, this included aligning the raw FASTQ reads to the hg38/GRCh38 human reference genome using BWA-mem 0·7·15,[16] sorting and removing duplicate reads with Picard 2·4·1. (http://broadinstitute.github.io/picard/), performing base quality score recalibration, indel realignment, generating single sample gVCFs with GATK HaplotypeCaller 3·5-0,[14] merging single sample gVCFs into batch specific VCFs (ranging in size from 40 to 588 quartets),[6] joint-calling the merged VCFs, and performing Variant Quality Score Recalibration (VQSR). The aligned CRAM and gVCF files were transferred to the Amazon Web Services (AWS) S3 storage system and can be accessed with permission from the Simons Foundation Autism Research Initiative (https://www.sfari.org/resource/sfari-base/).

The GS data from the fetal structural anomaly cohort was generated at the Broad Institute Genomics Platform. After sequencing, individual FASTQ files were transferred to a Google Cloud bucket for storage. All GS data pre-processing and sequence variant discovery was performed using the GATK Best Practices Workflows on the Terra platform.[17,18] Sequence variant calling followed the same steps described above for ASD.

1.2. Structural Variants (GATK-SV)

Structural variant (SV) discovery and genotyping was performed with GATK-SV, which was deployed deployment on the cloud-enabled and freely available Terra platform (https://terra.bio/). The code for GATK-SV is publicly available at https://github.com/broadinstitute/gatk-sv. All individuals were grouped into batches based on: 1) their dosage bias score (a metric that quantifies the non-uniformity of coverage for a given GS sample),[19] sex, family status, PCR status, and cohort assignment. The ASD cohort included batches comprising 200-400 samples each, and the fetal structural anomaly cohort included one batch of PCR plus (n=186) and two batches of PCR free (n=346 and n=346) samples, respectively. All families were kept intact during batching. All 7,195 individuals were analyzed with six SV discovery algorithms, including three paired-end/split-read algorithms (Manta v·1·4·0, Smoove v·0·2·3 [https://github.com/brentp/smoove], and WHAM-GRAPHENING v·1·7·0),[20–23] two read-depth algorithms (GATK-gCNV and cnMOPS v·1·12·0),[24–26] and one mobile element insertion algorithm, MELT v·2·0·5.[27] SV discovery generated six algorithm-specific VCFs per individual that were used as input for GATK-SV, which was run in cohort mode. The GATK-SV pipeline is organized into modules that harmonize predicted SVs across all input algorithms, reduce false positives, resolve overlapping SVs with disparate copy number, identify complex variants (*e.g.*, inversions flanked by one or more copy number variants [CNVs]),[28,29] and provide cohort-wide SV genotypes and quality metrics available for *post hoc* filtering. GATK-SV produced a cohort-wide SV VCF for the ASD and fetal anomaly cohorts, respectively, that was used as input for all downstream analyses. Further details on the GATK-SV methods can be found in Collins et al. 2020.[19]

1.3. Short tandem repeats (Expansion Hunter)

We identified short tandem repeat (STR) expansions across 18 loci that were selected from the gnomAD disease-associated STR catalog (https://github.com/broadinstitute/str-analysis/tree/main/str_analysis/variant_catalogs) based on conferring an early-onset developmental disorder phenotype (Table S6). STR expansions were genotyped using Expansion Hunter v5.0.0 across 6,435/6,448 individuals from the ASD quartet families (n=9 ASD probands with a sex chromosomal abnormality were removed as well as one quartet family that revoked consent after all other analyses were completed). We also applied Expansion Hunter to all 249 pre-screened fetal anomaly trios (n=747 individuals in total). Overall, Expansion Hunter identified 115,821 and 13,443 STR calls in the ASD and fetal structural anomaly cohort, respectively. Of these, 206 (0·18%; ASD) and 29 (0·22%; fetal structural anomaly) surpassed the literature-derived pathogenic repeat unit length and were visually assessed using REViewer.[30]

*2.0. Variant filtering*

2.1. Variant QC

After variant discovery, we applied quality control (QC) filters intended to maximize sensitivity for candidate P/LP variants while removing false variant calls. For SVs, this included removing variants with a GATK-SV QUAL score ≤ 1 and multiallelic copy number variants (CNVs). For sequence variants, we removed multiallelic variants, variants with an allele balance (AB) < 0.15 in the case of interest, indels > 50bp, and variants where the sum of the reference and alternate allele depth (AD) was ≤ 5 in any family member. We also removed SNVs that did not pass GATK VQSR. We also manually visualized read support for each candidate pathogenic STR expansion (n=235 across both cohorts) using REViewer,[30] which resulted in seven (five in the ASD cohort and two in the fetal structural anomaly cohort) candidates requiring manual variant interpretation. To reduce false positives, we applied additional quality control metrics to samples with outlier variant counts, defined as any sample with a variant count (based on raw GATK haplotype caller or individual SV algorithm output) above Q3 + 6*IQR. This definition resulted in relatively few SV outlier samples (n=12 SV in the fetal structural anomaly cohort and none in the ASD cohort) and sequence variant outliers (n=5 in the fetal structural anomaly cohort and n=1 in the ASD cohort). To control the false positive rate in these outlier samples, we removed SVs present in >2 SV outlier individuals and sequence variants with GQ <75.

2.2. Variant functional consequence

All variants (SNVs, indels, and SVs) were annotated for genic overlap and functional consequences against GENCODE v·26 gene boundaries.[31] SVs were annotated with GATK-SV, and all SV predicted to be loss-of-function (LoF) or full gene copy gain were retained for further filtering.[19] LoF SVs were defined as a deletion overlapping coding sequence, an inversion, mobile element insertion, complex SV, or translocation with one or more breakpoints disrupting coding sequence, or an intragenic exonic duplication (a duplication that overlaps coding sequence with both breakpoints contained within the same gene boundary). Full gene copy gains are defined as duplications that fully overlap a gene boundary. In this context, coding sequence refers to the longest confirmed protein-coding isoform from GENCODE v26. Partial gene duplications, defined as duplications with one breakpoint located outside the gene boundary and one within, were excluded given their unknown functional impact.[32] Sequence variants were annotated using ANNOVAR,[33] and any variants predicted to be stop-gain, stop-loss, frameshift insertion, frameshift deletion, splicing (within 2 bp of a splice junction [SO:0001568]), or missense according to RefSeq or Gencode annotations were retained for additional filtering. We further classified missense variants into three tiers (described below) to identify those that are increasingly likely to be functionally damaging and thus classified as P/LP:

**Tier 1 missense:**
- Missense variants classified as P/LP in ClinVar or with a CADD score[34] > 15
- Removed missense variants classified as benign, likely benign, risk factor, association, drug response, or protective in ClinVar

**Tier 2 missense:**
- Same filters as Tier 1, *plus*
- Missense variants with CADD scores between 15 and 30 had to be located in a missense constrained region[35]
- Any missense variant with a CADD score > 30

**Tier 3 missense:**
- Classified as P/LP in ClinVar

2.3. Disease genes and genomic regions

To facilitate variant filtration, we computationally built a candidate disease gene list for the ASD and fetal structural anomaly cohorts, respectively. The ASD gene list comprised 907 genes (Table S8) broadly associated with neurodevelopmental disorders (NDDs), including 902 genes from the DDG2P database[36] classified as having a 'confirmed' or 'probable' association with developmental disorders that conferred a brain/cognitive phenotype and 26 genes that were significantly enriched for rare *de novo* protein truncating variants in ASD (n=21 genes overlapped both lists).[4] To account for the variable phenotypes observed in the fetal anomaly cohort (Table S1), we compiled 2,535 developmental disorder genes (Table S3) based on the union of eight gene lists, described below:

1) 374 dominant developmental disorder genes from the DDG2P database (accessed July 29, 2019)[36] with a "confirmed" disease association and monoallelic, imprinted, mosaic, x-linked dominant, and x-linked over-dominance modes of inheritance.

2) 800 recessive developmental disorder genes from the DDG2P database[36] with a "confirmed" disease association and biallelic or hemizygous modes of inheritance.

3) 93 genes that were significantly enriched for rare *de novo* variants in the Deciphering Developmental Disorders study.[37]

4) 26 dominant genes significantly enriched for rare *de novo* protein-truncating variants in ASD.[4]

5) 358 genes from the Clinical Genome (ClinGen) Resource Dosage Sensitivity Map with "some evidence for dosage pathogenicity" (haploinsufficiency/triplosensitivity score = 2) or "sufficient evidence for dosage pathogenicity" (haploinsufficiency/triplosensitivity score = 3) (downloaded July 29, 2019; https://www.clinicalgenome.org/curation-activities/dosage-sensitivity/).

6) 708 autosomal dominant and 1,182 recessive disease genes curated from the Online Mendelian Inheritance in Man (OMIM) database.[38,39]

7) 217 recessive and dominant X-linked genes from OMIM (tables were accessed June 12, 2017).

8) 117 genes that have been robustly associated with fetal structural anomalies detectable by ultrasound that were curated by the Prenatal Assessment of Genomes and Exomes study.[40]

Each gene was classified as being associated with a disorder that had a dominant and/or recessive pattern of inheritance based on existing annotations from DDG2P and OMIM. We categorized the inheritance labels provided by DDG2P as recessive: biallelic, and hemizygous and dominant: imprinted, monoallelic, mosaic, x-linked dominant, and x-linked over dominant. When disease inheritance was not available for a gene (n=4 missing from DDG2P), variants in that gene were retained under both dominant and recessive modes of inheritance.

We also compiled a list of 65 known genomic disorder (GD) loci to assess overlap with SVs in both our cohorts. We took all of the known CNV syndromes located on the autosomes and chromosome X from DECIPHER[41] and the haploinsufficient (HI) and triplosensitive (TS) regions from the Clinical Genome (ClinGen) Resource Dosage Sensitivity Map if they had a HI or TS score ≥ 2 ("sufficient evidence for dosage pathogenicity"). We removed any regions that were only associated with late-onset conditions, resulting in 64 candidate regions (Table S4). All SVs that overlapped ≥ 50% of a GD locus were retained for manual review. Following the most recent guidelines for CNV interpretation,[32] we also manually reviewed any rare (<1% frequency in the genome aggregation database called gnomAD-SV)[19] deletion or duplication that overlapped ≥ 25 or ≥ 35 protein-coding genes, respectively, even if it did not overlap a disease gene or GD region from our lists. Finally, we also retained all SVs that overlapped one of 17 non-coding loci known to confer pathogenic long-range position effects (LRPEs; Table S5).

To define the non-coding search space, we used topologically-associated domain (TAD) boundaries from the IMR90 fetal fibroblast cell line,[42] which have been previously shown to be associated with pathogenic LRPEs if disrupted,[43–46] that contained each LRPE target gene.

2.4. Inheritance

We filtered variants under the five inheritance modes described below. For the ASD quartets, the unaffected sibling and both parents were treated as independent trios during inheritance filtering. We applied more stringent missense variant filters (tiers described in the variant functional consequence section of the appendix) to rare inherited and compound heterozygous variants as these two categories of variants have not been shown to substantially contribute to the aetiology of ASD or fetal structural anomalies.[2,4,40,47] The specific functional consequence considered for each inheritance type are as follows:

**Dominant disease genes:**

- *De novo*
    - All LoF
    - Missense Tier 1

- Rare inherited
    - All LoF
    - Missense Tier 3

**Recessive disease genes:**

- Homozygous
    - All LoF
    - Missense Tier 1

- X-linked recessive
    - All LoF
    - Missense Tier 1

- Compound heterozygous
    - At least one variant in the pair had to be LoF or Tier 2 missense

The identification of compound heterozygous variants comprised three steps, including: 1) compiling heterozygous SNVs, indels, and LoF SVs located in the same recessive disease gene, 2) annotating each variant with inheritance status, and 3) retaining only the instances where individuals had more than one variant in a recessive disease gene with disparate inheritance patterns (*e.g.*, one maternally inherited, one *de novo*). We required that at least one variant per compound heterozygous grouping be inherited from a parent due to the lack of phasing information from short-read GS and the high probability that at least one of the *de novo* events would be a false-positive or both would occur in *cis*.

2.5. Allele frequency

All variants (SNVs, indels, and SVs) meeting the above thresholds were retained if they had an alternate allele frequency (AF) <1% for dominant disease genes or regions and <5% for recessive disease genes. For sequence variants, this threshold was based on the maximum AF across gnomAD genomes, gnomAD exomes, ExAC, 1000 genomes, and the parental samples for each cohort.[9,48,49] SV AF was calculated based on the frequency of the event in gnomad-SV.[19] Given that some GDs can occur at an appreciable frequency in disease cohorts,[50] we did not apply any AF cut-off when considering SV that overlapped ≥50% of a known GD locus.

*3.0. Variant interpretation*

Details describing the methods for manual variant curation are described in the methods of the main text.

**Single sample SV pipeline**

We ran our single sample GS SV detection pipeline on all ASD probands identified to have a P/LP SV (n=77; 4.8%) to determine whether the cohort-based genomic analyses provided additional value and if GS-based SV discovery can be performed on a single sample with high sensitivity. To identify SVs in a single individual rather than across a cohort, we applied our cloud-based pipeline, which can be deployed through a publicly available workspace on Terra.[51] The stages of processing in the single sample pipeline are broadly similar to the steps described for the cohort-mode version of GATK-SV described above, with several notable differences: First, the GATK-SV single sample uses data and SV calls from a reference panel. In this study, our reference panel included 156 high-coverage GS samples that were randomly selected from the 1000 Genomes project.[52] We included an equal number of males and females, removed related individuals and samples with rare sex chromosome aneuploidies or evidence of mosaic chromosomal anueploidies, samples that had low absolute coverage, or were in the tails of the dosage score distribution[19] for the 1000 Genomes cohort. Second, portions of the evidence collection and variant filtering were replaced with the application of pre-trained models that were created by running the cohort mode pipeline on the reference panel. Third, variant quality filtering is replaced by a set of filters which examine the predicted genotypes of the reference panel samples for novel events detected in the case sample. SVs from single sample and cohort GATK-SV were considered the same event if they shared copy number, SV type, and had 75% reciprocal overlap of genomic coordinates.

**Benchmarking the performance of GS against conventional tests**

*Filtering CMA data*

As previously described,[5,12] SNP genotyping data was generated for the ASD cases using three microarray platforms, the Illumina 1Mv1, 1Mv3, or Omni2·5. CNV calls for each individual were identified using PennCNV,[53] QuantiSNPv2·3,[54] and GNOSIS.[55] CNVs were filtered for rarity based on overlap with CNVs from the Database of Genomic Variants (in GRCh36/hg18) and overlap with CNVs from the ASD parents.[12,56] All CNV coordinates were lifted over from GRCh36/hg18 to GRCh38/hg38 and those classified as high-quality (CNV p-value [pCNV] ≤ $1{\cdot}0 \times 10^{-9}$)[12] were filtered following the same steps outlined in the GS SV pipeline (Figure 1). There were 14 variants detected by GS that were also detected by CMA but failed filtering because they were not lifted over from hg18 to hg38 (n=6), failed the pCNV high-quality filter (n=5), or were removed due to incorrect CNV coordinates that suggested the variant did not overlap coding sequence (n=3; GS coordinates were used as truth). These variants were recovered and counted towards the overall yield of CMA. We also removed one deletion from CMA manual review that was identified to be rare by CMA but was found in 65 (2·0%) of our 3,224 ASD parents based on GS, which was above our allele frequency threshold.

*Filtering exome sequencing data*

The ES data for the ASD cases was generated as part of a larger sequencing initiative and has been extensively described.[4,12] We realigned sequencing data from GRCh37/hg19 to GRCh38/hg38[26] and applied the same filtering steps as those outlined in the GS filtering pipeline (Figure 1) with minor modifications to account for differences in depth between ES and GS (Figure S6). These included increasing our AB and sum(AD) filters for *de novo* variants (both in *de novo* dominant inheritance and as part of a compound heterozygous pair) to account for the higher ES coverage, increased rate of false positives, and potential for somatic variant detection. The new thresholds (sum[AD] ≥10 and AB >0·25) were chosen based on retaining >95% of the variants that were also detected by GS (Figure S6).

To identify CNVs from ES data, we applied GATK gCNV,[57] a publicly available Bayesian model for germline detection of CNVs.[57] Briefly, this is a read-depth based tool that uses a negative-binomial factor analysis to adjust for known and unknown biases of exome sequencing, while modeling sample and genomic region copy number through a hierarchical hidden Markov model. In this analysis, we jointly processed the 6,448 individuals from 1,612 ASD quartet families described in this study with an additional 66,000 samples.[26] Samples were assigned to batches based on 3D clustering of the first three principal components of coverage depth after normalizing for average depth. The 72,448 samples were processed across 126 batches, with a median batch size of 449 samples (min 136 and maximum 2,259). After raw calling with GATK gCNV, we applied our calibrated sample-level quality filters, resulting in 5·49% of the total samples being removed. The GATK gCNV quality score statistic (QS>400 for homozygous deletions, QS>100 for heterozygous deletions, and QS>50 for duplications) was applied to individual calls to extract rare CNVs with predicted sensitivity and positive predictive value of >90%, which resulted in a resolution of three exons or more. With these filtering metrics, the average ES sample harbored 1-2 rare high-quality CNVs.

#### SUPPLEMENTARY FIGURES

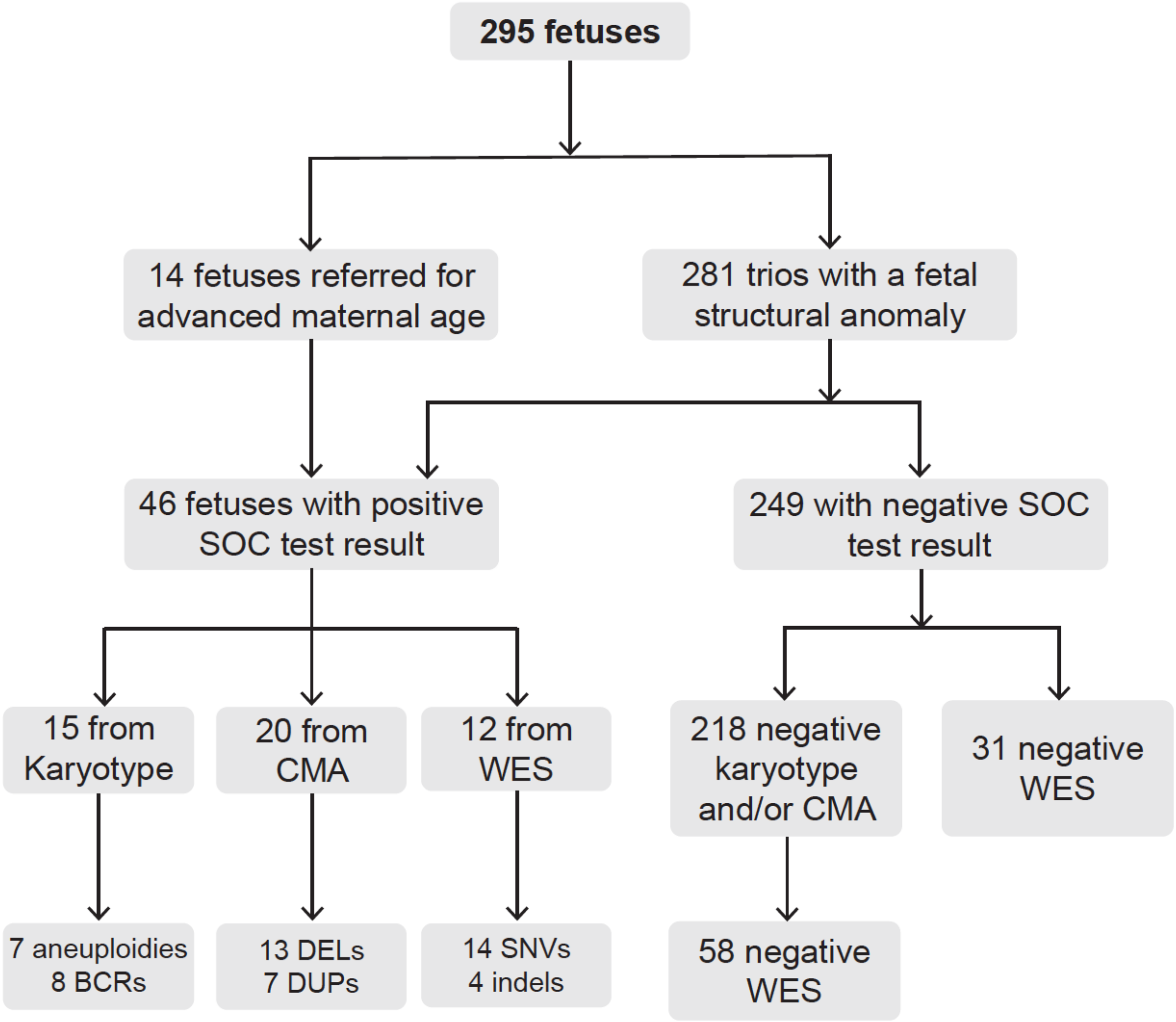


**Figure S1. Fetal structural anomaly study design**
Description of the 295 fetuses with GS included in this study, including the previous standard-of-care (SOC) diagnostic test results available for each fetus. CMA: chromosomal microarray, ES: exome sequencing, BCR: balanced chromosomal rearrangement, SNV: single nucleotide variant, indel: small insertion or deletion, Negative: no pathogenic or likely pathogenic variant identified, Positive: pathogenic or likely pathogenic variant identified.

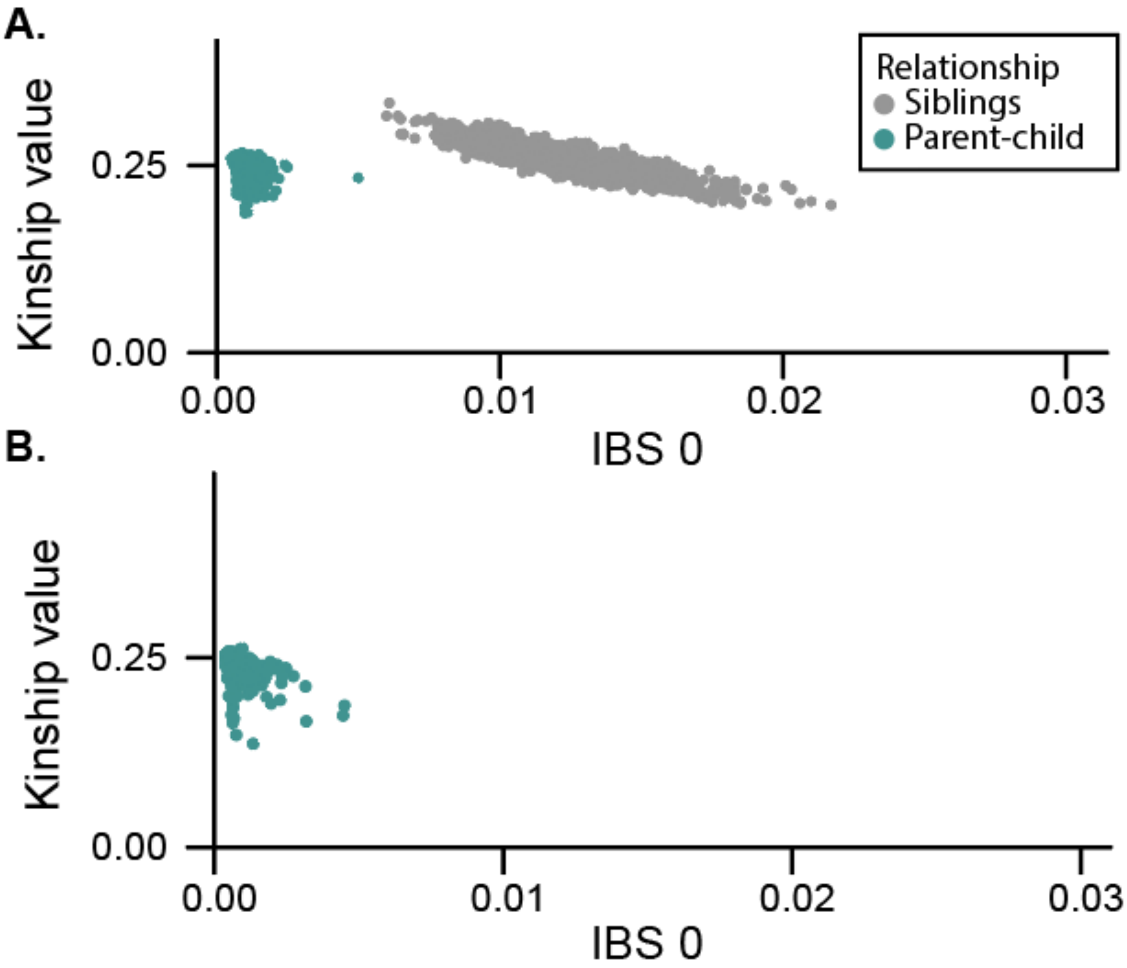


**Figure S2. Confirmation of sample relatedness using kinship values**
Kinship values for GS were calculated using KING[22] after restricting to SNVs with an alternate allele frequency greater than 5% in gnomAD genomes.[23] Each point on the plot represents a related pair of individuals, colored by relationship status. **(A)** Relatedness metrics for 6,448 individuals from the 1,612 ASD quartet families. **(B)** Relatedness metrics for 747 individuals from the 249 fetal anomaly trios.

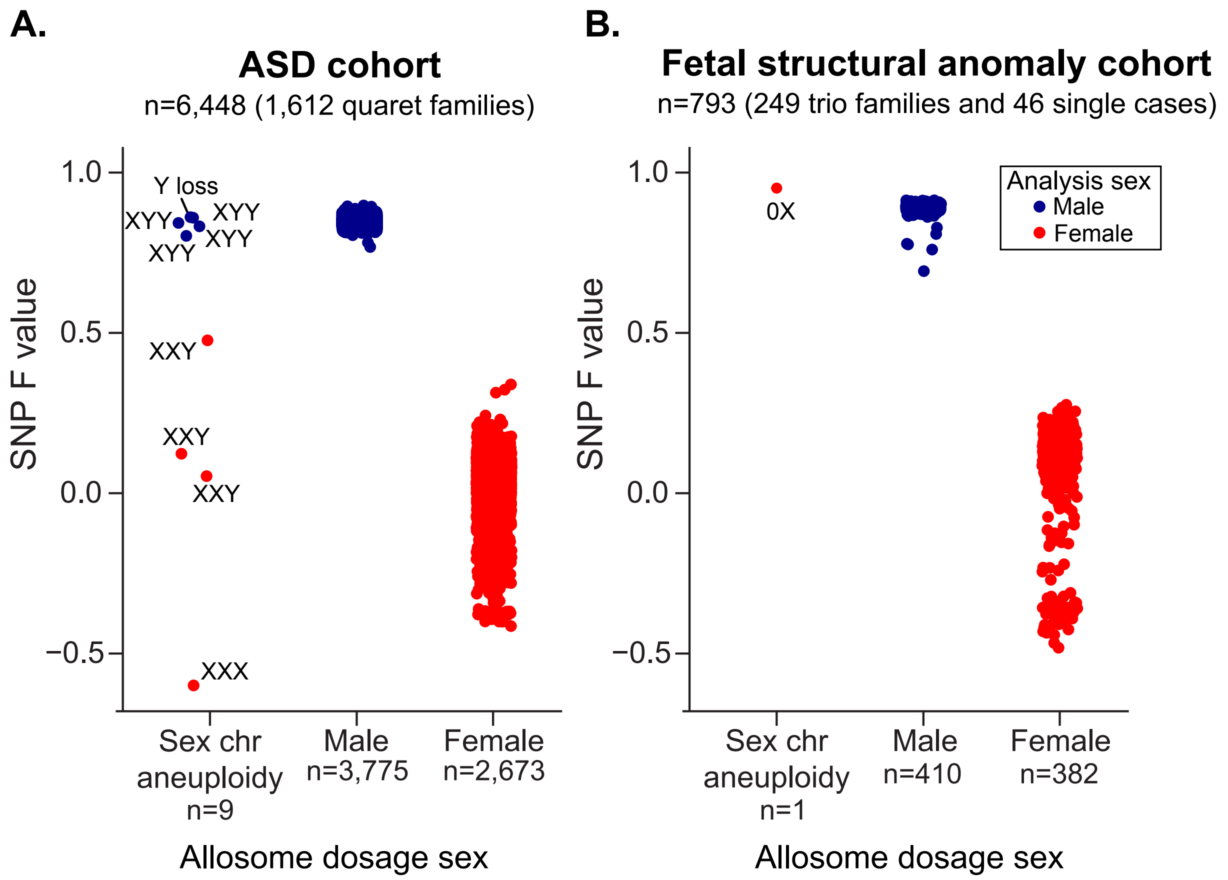


**Figure S3. Sample sex QC**

Confirmation of sample sex using single nucleotide polymorphism (SNP) and chromosomal read-depth information from GS data. Sex was inferred two ways from GS data: 1) using the F value generated with PLINK[24] based on sex chromosome SNP genotypes, and 2) using read depth (dosage) scores[19] derived from chrX and chrY. Each point represents a sample, colored by final sex used for analysis. **(A)** Sex metrics for the 6,448 individuals from the 1,612 ASD quartet families. Cases with sex chromosomal abnormalities (n=9) have been labelled. **(B)** Sex metrics for the 249 trios individuals from the fetal anomaly trios (n=747) that were pre-screened with standard-of-care diagnostic tests and the 46 single benchmarking cases.

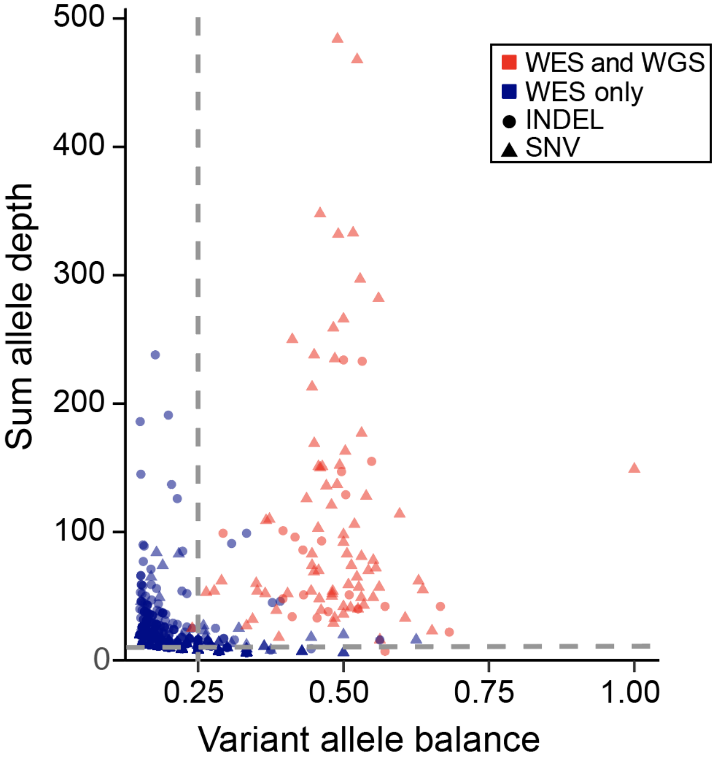


**Figure S4. Modified exome sequencing depth and allele balance thresholds**
The plot displays the allele balance (AB) and sum allele depth (AD) for all 1,453 *de novo* SNVs and indels detected from ES using the standard GS filters. Color indicates if the variant is unique to ES (blue; n=1,353) or was found in both the ES and GS data (red; n=100). Shape indicates variant type (SNV or indel). The dotted line represents the modified thresholds ultimately used for filtering the ES data before manual review: AB > 0·25 and sum(AD) ≥ 1.

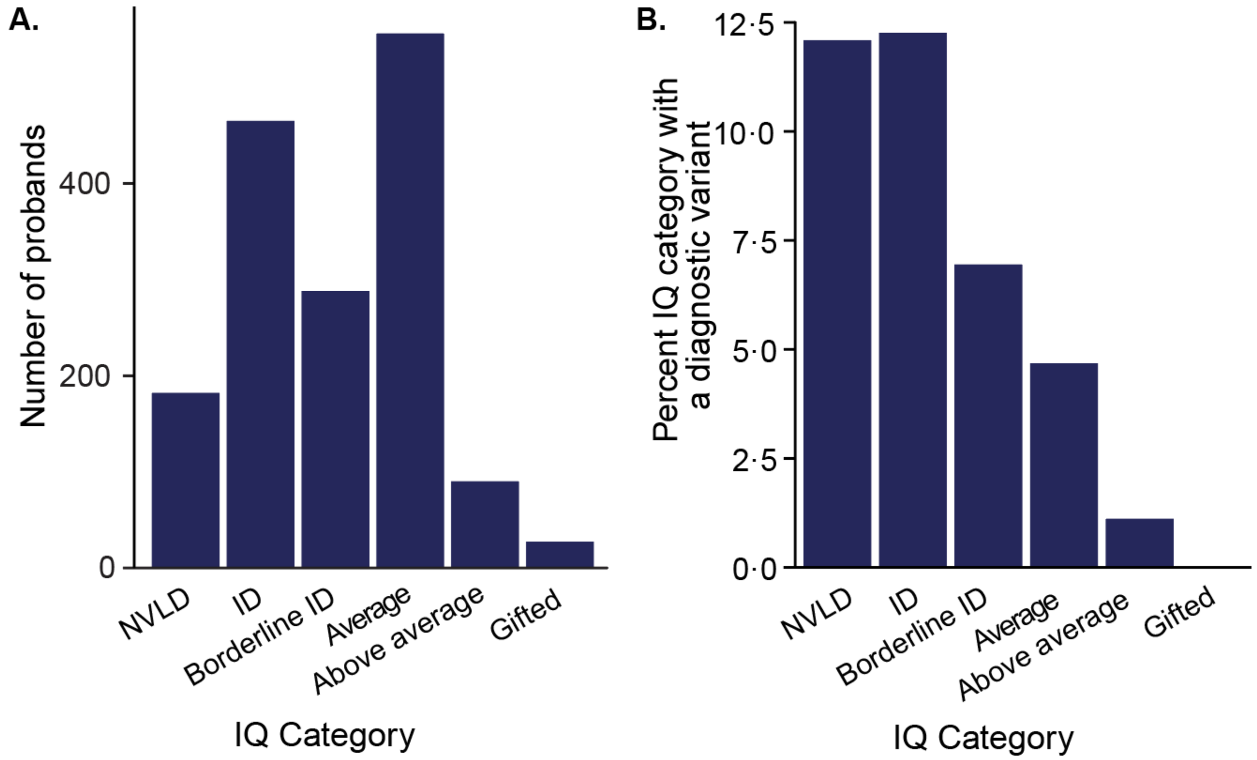


**Figure S5. Fraction of P/LP variants across IQ subgroups in ASD probands**
**(A).** Full scale IQ (FSIQ) for 1,608 probands in the ASD cohort according to the following IQ categories: NVLD: non-verbal learning disorder, ID: intellectual disability (FSIQ≤70), borderline ID (FSIQ 71-85), average (FSIQ 86-115), above average (FSIQ 116-130), and gifted (FSIQ>130). **(B).** Percentage of probands in each IQ bin that had a diagnostic variant (pathogenic or likely pathogenic) identified from GS analysis.

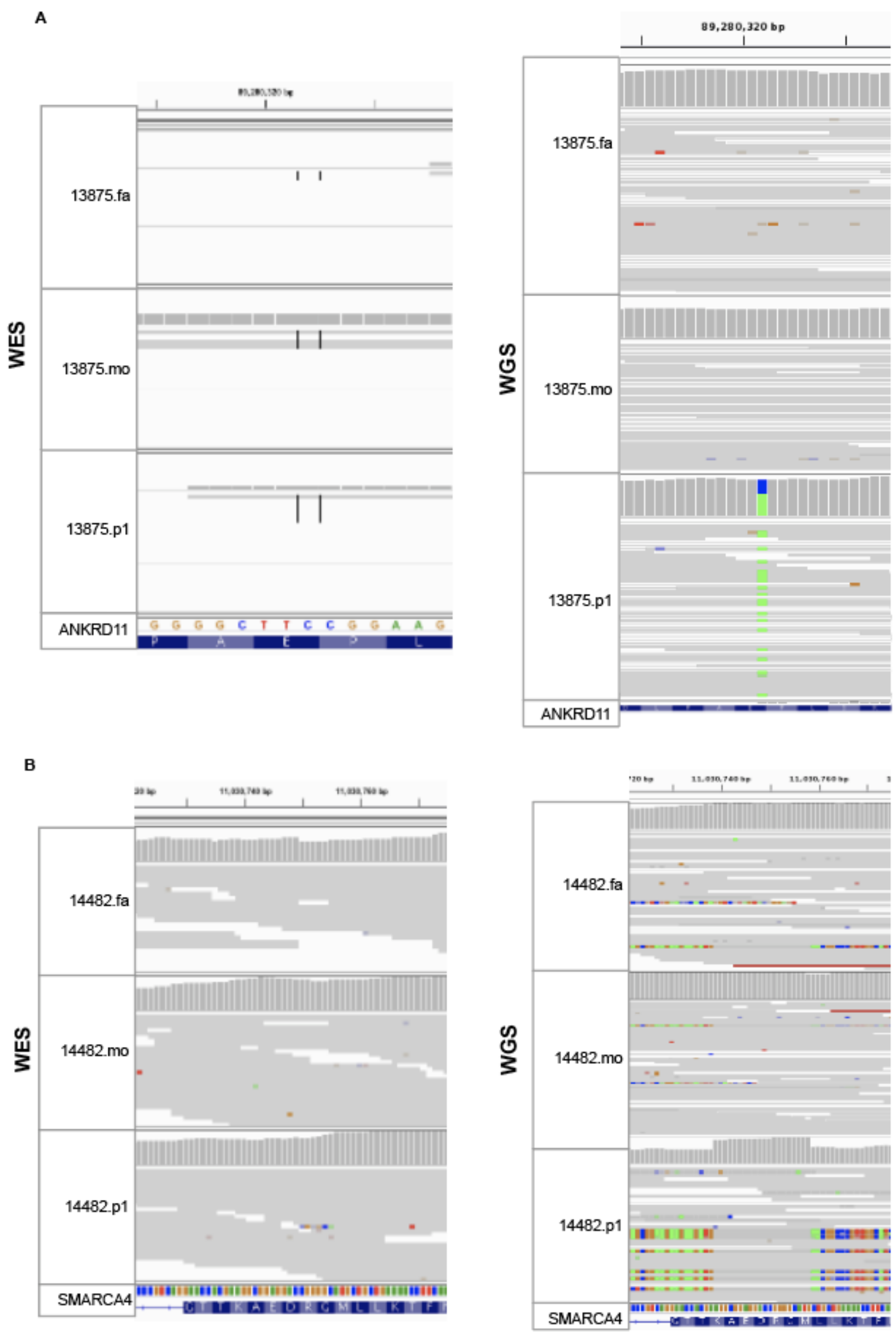


**Figure S6. Two pathogenic sequence variants unique to GS in ASD probands**
Alignment visualization for two *de novo* variants uniquely identified in GS alongside their raw read evidence from ES. Images were generated using IGV. For each site, the ES and GS screenshots are shown side-by-side. **(A)** A stopgain SNV in *ANKRD11* in proband 13875·p1 that was absent from the ES VCF and has low coverage in the ES CRAM files **(B)** A 44 bp insertion in *SMARCA4* in 14482·p1 that was absent from the ES VCF and shows no supporting evidence in ES CRAM files.

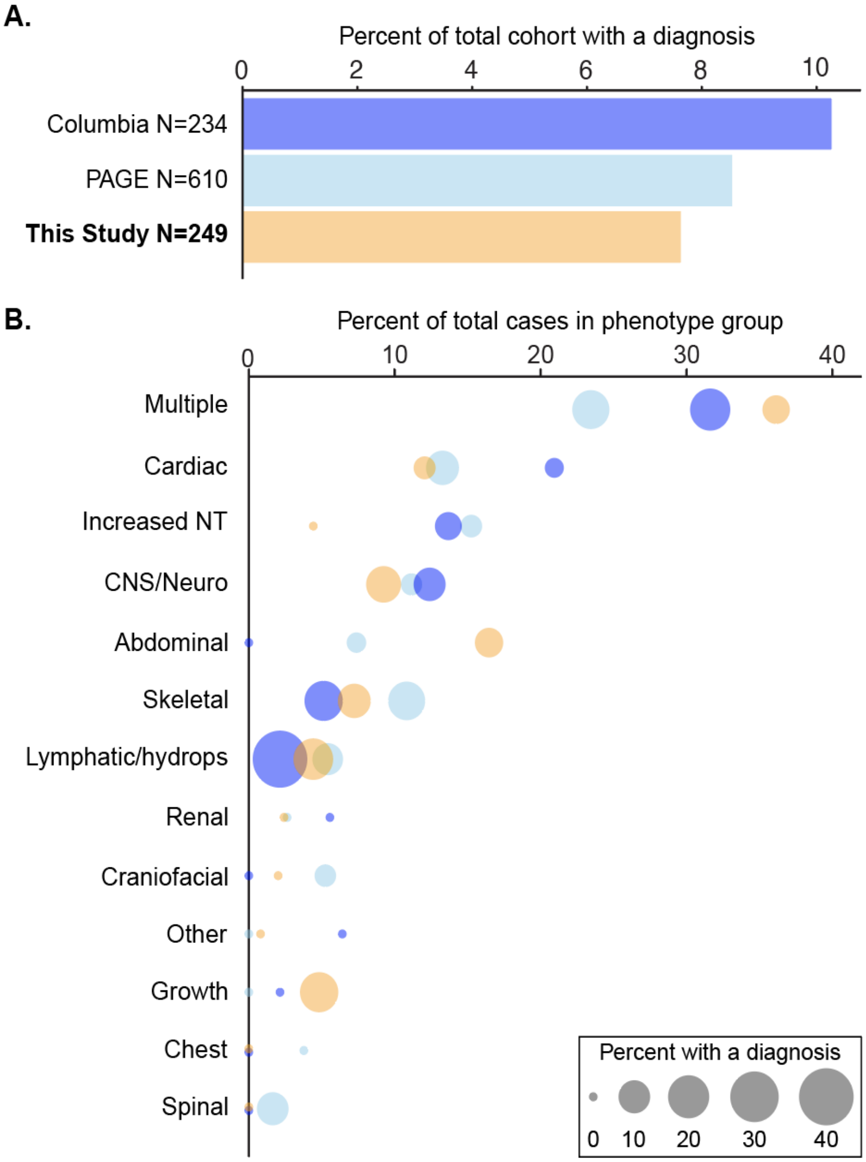


**Figure S7. Diagnostic yield of ES and GS across organ systems impacted by fetal structural anomalies**

**(A)** The fraction of P/LP variants identified by ES[2,40] or GS across three cohorts comprising fetuses with structural anomalies that were pre-screened using various standard-of-care diagnostic tests. **(B)** The number of fetuses with structural anomalies impacting each body system. Fetuses with more than one body system affected were considered to have "Multiple" anomalies and all remaining categories represented isolated anomalies. The x-axis represents the fraction of the total fetal cohort with an anomaly impacting each body system, the size of each point indicates the percentage of total cases identified to have a diagnostic variant within that category, and the color of the point corresponds to the study where the data originated.
